# Amazon biodiversity is at risk from metal contamination due to mining activity

**DOI:** 10.1101/2025.05.21.654597

**Authors:** Gabriel M. Moulatlet, Mariana V. Capparelli, Chris Thomas, Brad Boyle, Xiao Feng, Amy E. Frazier, César Hinojo-Hinojo, Juliana Herrera-Pérez, Lia N. Kajiki, Alex M. Lechner, Brian Maitner, Erica A. Newman, Efthymios I. Nikolopoulos, Patrick R. Roehrdanz, Lei Song, Daniel Valencia-Rodríguez, Wenxin Yang, Cory Merow, Miles Silman, Fabricio Villalobos, Mark Macklin, Brian J. Enquist

**Affiliations:** Department of Ecology and Evolutionary Biology; University of Arizona, Tucson, USA; Estación El Carmen, Instituto de Ciencias del Mar y Limnología, Universidad Nacional Autónoma de México; Carretera Carmen-Puerto Real km 9.5, 24157 Ciudad del Carmen, México; University of Lincoln; Lincoln, UK; Water & Planetary Health Analytics (WAPHA); Department of Biology, University of North Carolina; Chapel Hill, NC 27599, USA; Department of Geography, University of California; Santa Barbara, Santa Barbara California, USA; Departamento de Investigaciones Científicas y Tecnológicas (DICTUS); Universidad de Sonora, Hermosillo, Sonora, México; Laboratorio de Macroecología Evolutiva, Red de Biología Evolutiva, Instituto de Ecología A.C.; Veracruz, México; Amazon Charitable Trust; 42 Berkeley Square, W1J 6AW London, UK; Monash University Indonesia; Tangerang, Indonesia; Department of Integrative Biology, University of South Florida; St. Petersburg, FL, USA; Department of Biology, James Madison University; Harrisonburg, VA, USA; Department of Civil and Environmental Engineering, Rutgers University; New Brunswick, NJ, USA; Moore Center for Science, Conservation International; Arlington, VA, USA; Eversource Energy Center and Department of Ecology and Evolutionary Biology; University of Connecticut, USA; Sabin Center for Environment and Sustainability and Department of Biology, Wake Forest University; Winston-Salem, NC 27109 USA; The Santa Fe Institute; 1399 Hyde Park Rd, Santa Fe, NM 87501, USA

## Abstract

The Amazon basin hosts the most biodiverse and intact ecosystems on Earth, yet human activities are an increasing threat. Metal contamination due to mining constitutes one of these major threats, but its impacts remain poorly quantified. We provide the first quantitative assessment of biodiversity exposure to mining-associated metals—mercury [Hg], arsenic [As], copper [Cu], Zinc [Zn], and lead [Pb]—across the Amazon. Around 66% of the Amazon’s 38,890 species of birds, plants, mammals, reptiles, amphibians, and fishes are exposed to metal contamination, including biodiversity hotspots and Indigenous territories. Safeguarding the Amazon’s role as a global reservoir of biodiversity, ecosystem function, and cultural heritage requires addressing metal contamination not only as a localized issue, but as a pervasive threat to global biodiversity.

## Introduction

The Amazon basin is often regarded as one of Earth’s last ecosystems where ecological and evolutionary processes operate relatively undisturbed by industrial activity (*1*, *2*). This perception has shaped global discourse around conservation priorities (*3*). Yet, while deforestation, fire, and climate change dominate discussions of threats to Amazonian biodiversity (*4*), the expanding footprint of industrial and artisanal mining has received comparatively less attention. Metal contamination from mining, an often invisible but persistent form of environmental change, now pervades some of the most species-rich Amazonian ecosystems and river systems (*5*), driven in part by the intensified extraction of critical minerals such as mercury for gold and, increasingly, copper for the global energy transition (*6*)

Although environmental contamination is now recognized as one of five major drivers of global biodiversity loss (*7–9*), its spatial extent and ecological impacts remain unclear and poorly quantified, particularly in tropical ecosystems (*10*). Contaminants are potentially toxic elements of concern due to their persistence, toxicity, bioaccumulation, and biomagnification in high concentrations, posing risks to the ecosystem and human health (*11*). The gap in contaminant quantification reflects a broader bias in biodiversity science, where the contaminants are seldom integrated alongside land-use change, climate, and invasive species (*12*). Further, monitoring systems disproportionately focus on temperate, high-income regions (*13*). Developing spatially explicit approaches to assess large-scale contamination exposure is thus critical for understanding its ecological consequences on biodiversity-rich regions.

Mining in the Amazon basin predates European colonization (*14*, *15*), but over the last century, and especially the last decade, increasing demand for gold, copper, and tin dramatically intensified mineral extraction across the basin (*16–18*). More than 90% of metal contaminants from mining are transported via sediments, dispersing 10-100 km downstream, deposited and stored along river channels and accumulating in floodplains, where they persist for centuries (*19*). These sediments are continually remobilized by floods and fluvial dynamics, prolonging their ecological impact (*19*). Over time, these contaminants can then be assimilated into vegetation (*20*) and bioaccumulate in both aquatic and terrestrial animals (*21*), especially for higher trophic-level species (*22*), which can destabilize ecosystems and food webs (*23*). Disturbances such as fire can further mobilize these contaminants into the atmosphere (*20*) and disperse them into terrestrial and aquatic environments far from the original mining sources (*24*).

Metal contaminants impact biological processes ranging from cellular disruption to ecosystem dysfunction (*25–27*). Exposure pathways are diverse, affecting metabolism, growth, reproduction, species interactions, and community dynamics (*28*). Even sublethal contamination within a species’ geographic range may lead to immediate physiological effects and long-term population declines (*29*), cascading effects on trophic networks, and impacts on ecosystem stability (*22*). Given the broad and overlapping distributions of many Amazonian species, evaluating metal contamination requires spatially explicit, multi-taxa assessments. Integrating the increasingly available geospatial biodiversity data (*30*), spatial contaminant extents, and hydrological data (*31*) offers a promising framework to quantify species- and ecosystem-level exposure at large scale (*12*). Ultimately, metal contamination can potentially affect humans, especially Indigenous communities, whose diet is traditionally composed of local fish and game (*32*). Quantifying the proportion of species’ geographic ranges intersecting contaminated areas is thus essential for anticipating poisoning risk and guiding conservation strategies.

Here, we address a critical gap in Amazon biodiversity science: the lack of spatially resolved assessments of species exposure to chemical contamination across taxonomic groups. Specifically, our analysis provides the first quantitative assessment of biodiversity-wide exposure to metal contamination across the Amazon, Earth’s most biodiverse terrestrial ecosystem, with significant reach across diversity hotspots and biogeographic provinces. Our framework offers a scalable approach to assess contaminant-driven risk in this and other highly biodiverse, metal-mined regions such as Southeast Asia and West Africa. Further, this approach underscores an urgent concern: as climate change accelerates species range shifts across Amazonia, the spatial overlap between biodiversity and contaminated areas will likely expand, compounding exposure to different metal types and amplifying the long-term ecological footprint of mining.

We combined a comprehensive database of species geographic ranges for 38,815 species (Figure 1A) of plants and five vertebrate groups (fishes, birds, reptiles, mammals, and amphibians; [(*33*, *34*)]) with recent maps of mining contamination. The maps of mining contamination in the aquatic ecosystems for As, Cu, Pb, and Zn come from (*19*), estimated as the attenuated dispersal of contaminated sediment downstream from mapped mine operations along the rivers, associated with formal sector mining of 21 metal mineral commodities. For Hg, we incorporated informal, artisanal mining data from Amazon Mining Watch (https://amazonminingwatch.org; accessed in October 2024) (Figure S1). To find the most contaminated areas, rivers reaches (500m) were considered contaminated if they were within the distance from a mining source with an associated contaminant metal concentration predicted to be above the Probable Effect Level (PEL [(*35*)] — the threshold above which adverse effects are frequently expected to occur in living organisms (Figure 1B) (*35*).

**Fig. 1.**
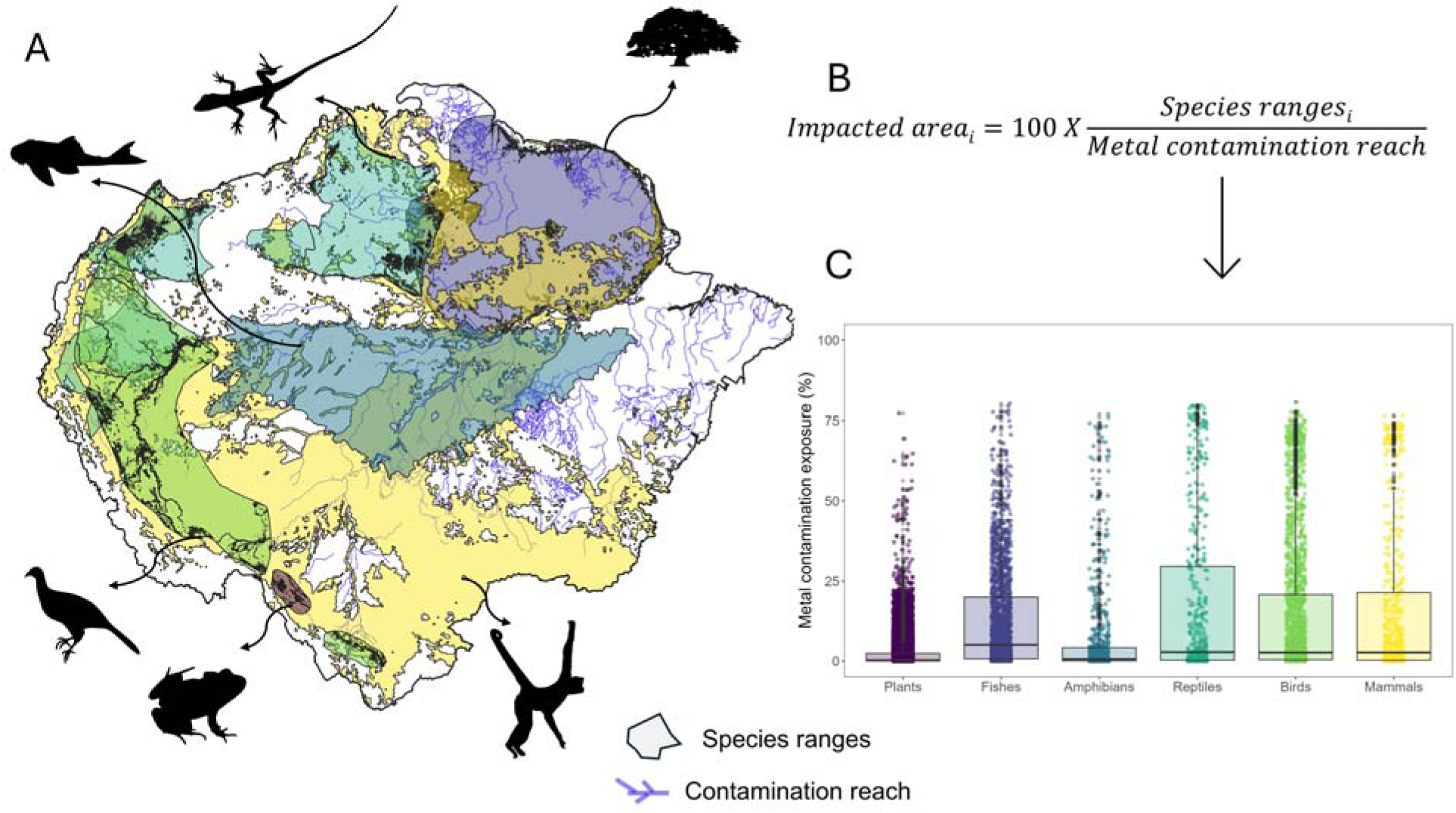
How to calculate the species’ geographic ranges exposure to metal contamination? **A)** Example of the geographic ranges for a single species of each taxonomic group in the Amazon basin. Blues lines in the background indicate the extension of a given metal along the rivers. **B)** For each species *i*, we calculated the proportion of its geographic distributions that overlaps with the rivers where metal contamination (either As, Cu, Hg, Pb or Zn) has been detected. **C)** Fractions of Amazonian species geographic distributions exposed to metal contamination sorted by taxonomic groups. We calculated the impact for 744 mammals, 1212 amphibians, 661 reptiles, 2,361 birds, 31,759 plants and 2,150 fish species. Average values are shown in Table S1 and individual species values in Appendix 1.

While our framework does not capture ecological amplifiers, such as bioaccumulation across trophic levels or bioavailability variation due to environmental interactions (e.g., Hg-organic carbon interactions), it identifies priority areas where mining impacts are most likely to threaten species. Therefore, results presented in this paper may be taken as lower range estimates on exposure. By mapping these exposures, we provide new information of the multiple threats to biodiversity, highlight urgent targets for mitigation and underscore the need to treat mining contamination in freshwater not as a localized issue, but as a pervasive threat to global ecosystems.

## Results

### Most Amazonian species are exposed to metal contamination

Our analysis shows that 5% of Amazonian aquatic streams and rivers (1,197,674 km^2^) had metal-contaminated areas (64,076 km^2^). Of the 38,887 assessed species, 66.52% (25,871) had ranges with metal-contaminated areas (Figure 1C). This includes 74% of birds, 68% of mammals, 60% of fishes, 52% of reptiles, 35% of plants, and 38% of amphibians (Figure S2). The fraction of a species’ geographic range exposed to metals varied across taxa (Appendix 1). On average, reptile, mammals, birds and fish species had the highest contamination exposure (17.8%, 15.9 14.6 and 12.4 of their geographic ranges intersect with metal reaches, respectively), while plant and amphibians had the lowest exposure (2.2% and 6.8% of their geographic ranges, respectively) (Table S1). Notably, 70 fish species (3.2% of all fishes), 88 reptiles (13.3% of all reptiles), 249 birds (10.5% of all bird species), 96 mammals (12.9 % of all mammals) and 15 plant species (0.05% of all plants) had more than half of their distributions exposed to mining-associated contamination (Appendix 1).

Exposure also varied among different metals. Mercury (Hg; 37,778 km^2^) and arsenic (As; 17,716 km^2^) exhibited the largest spatial footprints (Figure S1), being present in the most significant fractions of species’ ranges (Appendix 1). Our results indicate that, on average, 9.2% of all fish geographic ranges are directly exposed to Hg, while 3% of all species analyzed faced As, Pb and Cu exposure. Zn contaminated less than 1% of species’ ranges (Figure 2, Table S2). Moreover, more than 40% of all species of the different taxa are exposed to As and Hg and more than 30% to Cu and Pb (Table S2).

**Fig. 2.**
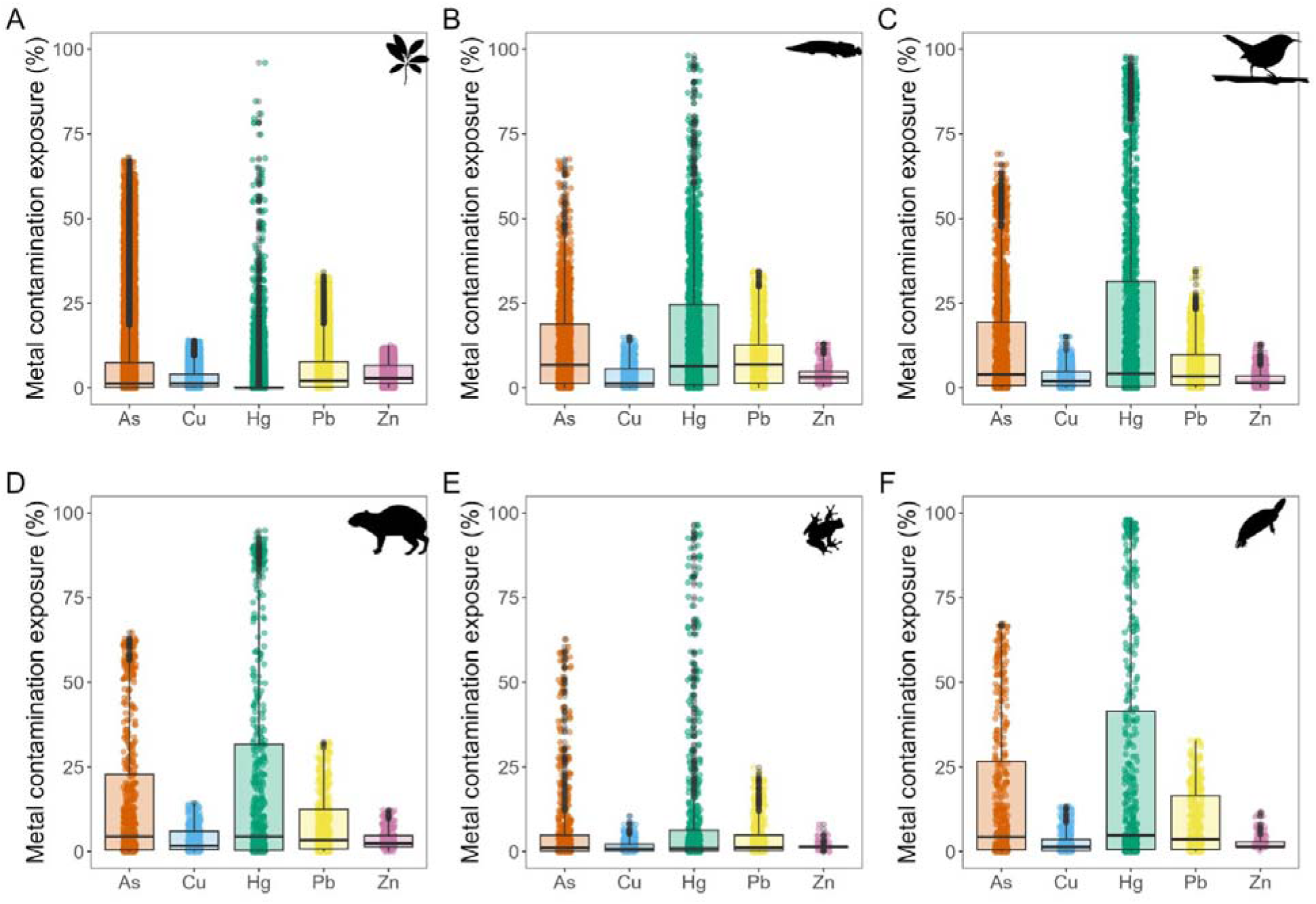
Fractions of species geographic ranges within the Amazon basin exposed to metal contamination derived from mining activities. Results are shown for **(A)** plants, **(B)** fishes, **(C)** birds**, (D)** mammals, **(E)** amphibians and **(F)** reptiles. Minimum, maximum and average values are shown in Table S2 and individual species values in Appendix 1.

### Plant species highly exposed to metal

From the plant list, 276 were identified as riparian tree species specialists (*36*). Unlike other *terra-firme* plant species, these riparian trees have geographic distributions that primarily encompass the flooding zones along Amazonian rivers (i.e., várzeas and igapós), where our models have mapped the majority of metal contamination. On average, riparian trees had 37% of their range exposed to metal contamination, approximately 18 times greater exposure than other plant species (2.2%). Mercury and arsenic were the primary contaminants, affecting 45% (40 times more than non-riparian plants) and 33.4% (6 times more than non-riparian plants) of exposed ranges respectively (Figure S3). Although the physiological effects of chronic metal exposure on Amazonian trees remain poorly understood, some riparian species are known metal bioaccumulators (*27*). Exposure to contaminated water and sediments can impair nutrient absorption, reduce growth rates, and impact survival (*37*). Moreover, plant species near artisanal gold mining intercepts large amounts of particulate and gaseous mercury, with substantial accumulation in biomass (*20*). Of particular concern are the edible fruits and seeds produced by many of these trees, which are consumed by wildlife, creating potential pathways for trophic transfer and human exposure to metals, ultimately affecting public health as well (*38*, *39*).

To assess additional to further cascading spread metal contaminants to the atmosphere we assessed fire as a threat multiplier. We used the results of (*40*) to identify 4,227 species (13% of all plant species) exposed to both fire and metals. Out of those species, 167 that have more than 5% of their geographic ranges exposed by both fire and metals and nine species exceeding 10% exposure for both threats (Appendix 2). Among the species with more than 10% of its range exposed to both threats are *Terminalia lucida* (Combretaceae), *Inga bullatorugosa* (Fabaceae), and *Ouratea polygyna* (Ochnaceae). These species are also listed among those that are not hyperdominants, i.e. those whose populations are composed of a relatively small number of individuals, and therefore, that may rapidly decline with increasing anthropogenic impacts (*41*).

### Fish exposed to metal contamination enter the food chain

Our analysis reveals considerable metal exposure among commercially important Amazonian fish species, with potential implications for human health and ecosystem functioning. Of the 2,150 fish species assessed, 10 are intensively commercialized (*42*), including *Arapaima gigas*, *Brachyplatystoma rousseauxii*, *Colossoma macropomum*, and *Zungaro zungaro* (Figure S4A), and showed substantial mean range overlaps of 19.7 % with contaminated areas, ∼2 times higher than for the other fish species. The exposures of commercial fish species were higher for 25% (Hg), 21% (As), and 15.5% (Pb). This result corroborates recent reports of metal contamination in Amazonian fish markets (*43*) and raises particular concern for methylmercury (MeHg), a potent neurotoxin that readily biomagnifies through food webs from low environmental concentrations (*44*). This is followed by biomagnification through successive trophic levels, ultimately resulting in the accumulation of MeHg in fish (*45*), representing risks to human health (*46*). Several fish species are under threat due to impacts caused by humans on the aquatic environments (*47*). Notably, species classified as Least Concern by the International Union for the Conservation of Nature (IUCN) red list criteria exhibited the highest proportional exposure to metals (Figure S4B), suggesting that threat assessments may underestimate contamination risks. While these species face fewer conservation pressures overall, metal contamination represents a growing threat to their populations.

### Diversity hotspots and Indigenous territories are highly exposed to metal contamination

By estimating species’ exposure to different metals across their geographic ranges, we determined the regions in Amazonia with the highest metal exposure, with eastern Amazonia showing the highest overall species exposure (Figure 3, Figure S5). Taxa exhibited distinct spatial patterns: plants had the most significant exposure to metals in central and eastern Amazonia (Figure 3A). Fish showed elevated exposure in eastern and western Amazonia (Figure 3B). Amphibians face great exposure in northern and eastern Amazonia (Figure 3C). Reptiles had the highest exposure in north and western Amazonia (Andes foothills) (Figure 3D); birds in central, north and eastern Amazonia (Figure 3E), and mammals in eastern Amazonia (Figure 3F). All these mapped areas have historically been mined for the valuable metals found along river terraces (such as in western Amazonia). Large-scale mining projects in these areas have left persistent metal loads exceeding PEL thresholds, and therefore, more species with impacted geographic ranges (*48*).

**Fig. 3.**
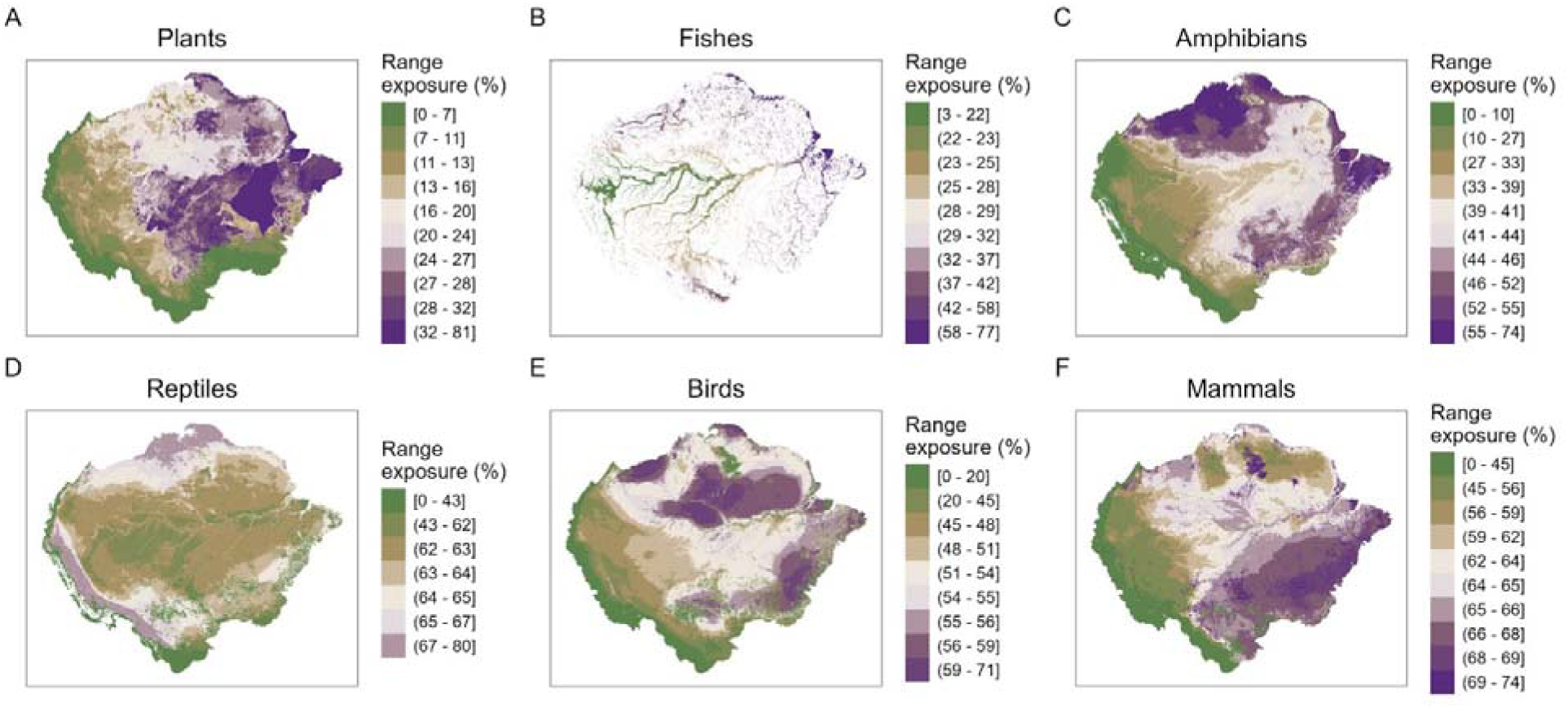
Map of species geographic distributions exposure for all species. Geographic distributions exposure maps were made by assigning the corresponding values to their exposed areas (one value for each species geographic range; Figure 1), and then calculating the median values per each 5 km by 5 km grid cell. Results are shown for **(A)** plants, **(B)** fishes, **(C)** birds, **(D)** mammals, **(E)** amphibians and **(F)** reptiles.

By overlapping species geographic ranges, we created species richness maps for Amazonia (Figure S6), identifying diversity hotspots as those areas where species richness was in the 50th, 25th, or 10th percentile or higher. The overlap of these hotspots with the metal contamination zones revealed the areas where biodiversity is more exposed to metal contamination (Figure S7, Table S3). For areas with species richness in the 10th-49^th^ percentile, all regions had less than 4% overlap with contaminated areas. The overlap with contaminated areas is considerably higher, at 11-14% for species richness at the 50th - 99th percentile in all regions, except for Eastern Amazonia, where the overlap is below 2% (Figure S7 A-E, Table S3).

The diversity hotspots that overlapped with contaminated areas varied for each taxon (Figure S8, Appendix 3). When diversity hotspots were defined as the top the 50th percentile of species richness by taxa, reptiles (26%), fishes (20%), amphibians (18%) and birds (16%) had larger exposure to metals in Southern Amazonia. In comparison, mammals (21%) and plants (14 %) had larger exposure areas in Northern Amazonia. Diversity hotspots exposed to metal contamination were higher in Southern and Central Amazonia, regardless of the chosen richness percentile thresholds (Figure S9). Amazonian ecosystems in these regions may have already approached their climatic and disturbance tipping points (*4*), impairing their ability to recover from a degraded state to a mature wet forest. Species inhabiting degraded ecosystems are more likely to face intensified adverse effects of metal contamination (*49*, *50*).

Indigenous territories are key areas for conservation of Amazonian human culture, ecosystems and protection of animal and plant species (*51*). Identifying the overlap of Indigenous territories with the areas where species are exposed to metal contamination can indirectly indicate the vulnerability of these human populations to contaminated food. Indeed, all the Indigenous territories in our dataset (*n* = 409) have species exposed to metal contaminations (Appendix 4). The highest exposure in an Indigenous territory was 64% of species, with a mean exposure of 35%. Our findings imply widespread risks to food security, as the presence of animals and plants exposed to metals in these territories may introduce metals into Indigenous diets (*52*, *53*)

## Discussion

### The exposure of Amazon biota to metal contamination is vastly underestimated

The metals evaluated in our study have been shown to play critical roles in metabolic processes at low concentrations, but toxicity varies with concentration (*54*) and by metal contaminants. While Cu and Zn are essential micronutrients for several species at low concentrations (*55*), our modeling focused on levels exceeding the probable effect level (*35*), suggesting that they are found at concentrations that generate harmful effects for the biota (*56*). Hg, As, and Pb are non-threshold toxins and are classified as potentially toxic metals at all concentrations, but the duration and magnitude of exposure determine the eventual outcomes. For Hg, As, and Pb, numerous fatalities in animals have been reported, particularly in blood composition, lung function, and brain activity, even at low doses (*57*). Arsenic is known to bioaccumulate in aquatic organisms. However, arsenic toxicity is highly dependent on its chemical speciation, and unlike Hg, and its potential biomagnification in food webs is low (*58*). Nevertheless, both metals bioaccumulate in terrestrial and aquatic ecosystems near historical gold mining sites (*45*).

Mercury contamination is of particular concern because, compared to the other metals mainly transported by water and deposited in the sediments, it has high volatility and is transported by air in the form of MeHg. Mercury compounds can become airborne during the burning of Hg-gold amalgam at or close to mining sites (*20*), resulting in mercury compounds becoming deposited as wet or dry deposition to foliar surfaces and soils (*20*, *59*). In Madre de Dios, Peru, gaseous elemental mercury was reported in concentrations 10 to 5000 times greater than the background concentrations in locations 100 km from the mining site (*60*). Moreover, gaseous elemental mercury was detected in the hair of Indigenous people 300 to 400 km upstream from mining sites in the Manu region, also in Peru (*61*). This exposure was associated with decreased IQ scores. Air transport is a significant pathway for Hg (*62*), but it is uncertain how that mode of exposure translates into input into aquatic systems. Accounting for gaseous elemental mercury in the soils would provide a complementary estimation for the exposure of several species used in this study, as we found that the exposure of fish species to Hg was ∼ 1000 times greater than for terrestrial plants and vertebrates.

Our analysis is based on contamination along mapped river networks with upstream catchments larger than 10 km², thus excluding smaller headwater streams that harbor ecologically important and biologically rich diversity (*5*, *63*). This exclusion likely underestimates contamination extent and species exposure, particularly given that aquatic environments comprise nearly 20% of Amazonian landscapes (*64*). During routine movements, terrestrial and semi-aquatic species frequently encounter smaller water bodies (i.e., small streams, ponds, or floodplains), potentially exposing them to contaminated sediments beyond our detection resolution. Furthermore, Amazonian rivers are highly dynamic and may change their course within less than a decade (*65*), altering contamination exposure patterns and biogeographic connectivity. Such hydrological changes may create new dispersal barriers while increasing species’ contact with contaminated zones (*65*). Continued monitoring is essential to account for these temporal shifts and their ecological consequences.

Several factors contribute to additional underestimation in our exposure assessments. First, we evaluated only the five major known metal contaminants in isolation, whereas natural systems typically contain metal mixtures with potential synergistic or cumulative effects (*66*). Second, our analysis excludes other mining-related impacts, such as deforestation, road openings, river channel interruption, and river diversion. Despite these limitations, our study represents the first attempt to map the extent of mining contamination across the geographic distributions of a large number of plant and animal species in the Amazon, establishing a crucial baseline for future research, policy, and conservation strategies. This urgency is further amplified by growing global demand for critical minerals, such as copper, and gold which is accelerating both legal and illegal extraction activities across the Amazon basin.

### Contamination is reshaping Amazon biogeography

The Amazon is widely believed to be one of the last remaining large ecological landscapes where biological and ecological processes continue largely unimpeded by industrial human activity (*2*), fostering the perception that much of the basin remains ecologically intact. Our findings challenge this assumption. Large areas of the basin—long considered to be beyond the reach of human contamination—are now contaminated by persistent and toxic mining-derived metals. These persistent contaminants pose growing risks to ecosystems and human populations, particularly rural and Indigenous communities who directly rely on rivers for cultural, spiritual, and subsistence needs (*67*, *68*). The threat extends beyond metals, as pharmaceuticals, pesticides, fertilizers and other contaminants from mining and urban expansion increasingly degrade Amazonian freshwater systems (*69*, *70*).

Mapping of species’ geographic ranges intersecting metal-contaminated rivers identifies areas of potential population-level risk. While not directly assessing toxicological impacts, we provide a spatial framework for identifying species and regions likely exposed to contamination levels above known effect thresholds. For instance, if a large portion of a species’ geographic range overlaps with metal-contaminated areas, this suggests widespread exposure that could impair populations’ viability and have a cumulative effect of metal accumulation across trophic levels. Conversely, minimal overlap indicates populations are likely to face lower contamination risk. As such, our findings can help target and prioritize future research efforts to better understand the impacts of metal contamination on all levels of biological organization, including individuals, populations, ecosystems, and extending to human health and culture.

Climate change will further complicate this picture by altering the geographic ranges of thousands of Amazonian species, as temperature and precipitation regimes change across elevational and latitudinal gradients (*71*, *72*). Species may shift their ranges to track suitable climate conditions, and many may encounter contaminated river corridors and floodplains, exposing previously unaffected populations to metals. This dynamic could simultaneously increase contamination exposure while constraining species’ future adaptive responses to climate change, reducing the viability of critical climate refugia. Metal contamination thus represents both an immediate threat and a future constraint on Amazonian biodiversity’s response to global change.

### Governance gaps and enforcement challenges in mining regulation

Current regulatory frameworks fail to control mining impacts across Amazonia adequately. For example, local authorities in the Amazon tasked with preventing and enforcing law around mining activities encounter several obstacles to achieving this goal (*5*, *73*), even though enforcement is an effective deterrent and governance tool (*74*). Access to mining sites is frequently restricted, rendering regulations unenforceable. Existing policies are often inadequate or not tailored to the ecological realities of Amazonia (*29*), and usually dismantled (*75*). Without regional-scale monitoring, the extent of biotic and abiotic impacts will remain difficult to anticipate or manage. Since mining occurs across all major sub-basins (*48*), bioaccumulation of metals in local biota is likely widespread (*21*, *76*). The spread and fate of contaminants in fluvial systems are complex and long-lasting (*29*). It is therefore critical to challenge the absence of a more severe law enforcement perspective related to opening new legal mining sites in the Amazon and the uncontrolled illegal mining activity (*77*). Our findings demonstrate that persistent mining-derived contamination is likely impacting many species, potentially reshaping the biogeography of the world’s most biodiverse forest. Moreover, our findings provide a baseline of species exposure to metal-derived contaminants that can be useful for policymakers and conservation. Many species affected - including plants and animals used for food and medicine - may already accumulate potentially toxic metals, with unknown implications for ecological and human health (*63*). To safeguard the Amazon’s role as a global reservoir of biodiversity and ecosystem function, metal contamination from mining must be recognized not as a localized issue, but as a pervasive ecosystem change driver (*3*). Therefore, metal contamination should be treated as a widespread problem critical to impact assessments and conservation planning.

## Materials and Methods

### Species geographic ranges

We obtained the species ranges from several sources. For fish, we first selected species with available geographic range maps (*78*), resulting in 2,442 species with available distributional data in the Amazon basin, representing 97% of all fish species reported for the basin (*79*). Then, we obtained their geographical ranges (extents of occurrence) from a previous study (*78*). There, the geographic ranges of freshwater fish species were constructed by creating a convex hull for each species based on its occurrence records and overlapping it with the HydroBASINS layer level 8 (*80*) from the HydroSHEDS database (https://www.hydrosheds.org/) to obtain geographic ranges in sub-basin units. Out of those 2,442 species, we identified species of commercial interest based on the list provided by (*42*). Threatened category data for fish species was reviewed and downloaded from https://www.iucnredlist.org in November 2024.

For mammals and birds, geographic ranges were built using Area of Habitat (AOH) maps downloaded directly from (*34*). For migratory birds, separate AOHs were generated for the resident, breeding, and non-breeding geographic ranges, corresponding to the species’ migratory distribution. For reptiles and amphibians, we built geographic ranges using the AOH workflow (*34*, *81*) with the R package “rgrass” (*82*). The AOHs are the remaining areas after removing those with known unsuitable habitat types or beyond elevation limits within each species’ geographic ranges. We collected expert-based geographic range maps, habitat type, and elevation information for each species from the IUCN Red List. We first excluded species without habitat information, whose habitat code was exclusively marine, subterranean, or cave, or whose IUCN category was extinct or extinct in the wild. We retained geographic ranges where species presence was extant or probably extant, and species origin was native, reintroduced, or present due to assisted colonization. We then created a base map to mask areas beyond species elevation range and suitable habitat types based on a translation matrix between habitat types and land cover types (*81*). The base map was created by combining Copernicus Global Land Service (CGLS) Land Cover data (*83*) and NASA Shuttle Radar Topography Mission (SRTM) elevation data (*84*), and each pixel’s value was equal to 1000 times land cover code plus elevation divided by 10. The CGLS Land Cover data were produced at a 100 m resolution, and the SRTM data were resampled to match this resolution. Species whose geographic ranges were exclusively in islands not covered by the land cover map were not included.

For the plant species, we used a previously validated workflow (*33*) with the BIEN (Botanical Information and Ecology Network) database (*40*) to obtain species occurrence points and to construct geographic ranges for each species. In brief, we used species distribution models provided in BIEN to infer the geographic distributions of species. These distribution models are based on occurrences from the BIEN database (Botanical Information and Ecology Network v4.2.8; http://bien.nceas.ucsb.edu/bien/about/) and climate and soil characteristics described below. BIEN occurrence records are compiled via a linked workflow that standardizes, integrates, corrects and validates data from disparate data sources and formats, including herbarium collections, ecological plots and surveys, and trait observations from various sources (Supplementary Information).

Extensive data cleaning was performed to ensure the reliability and harmonization of records compiled from different data contributors to the BIEN database. Taxon names associated with BIEN occurrence records were corrected and standardized using the Taxonomic Name Resolution Service (TNRS) (*85*). When available in the original data, family information was included along with the species name submitted to the TNRS. This information enables the TNRS to detect homonyms (identical names referring to different taxa) in other families. The declared political division names of occurrences were standardized using the Geographic Name Resolution Service (GNRS; http://bien.nceas.ucsb.edu/bien/tools/gnrs/) (*86*), which corrects spelling errors and standardizes names, codes, and abbreviations. Occurrence records that fall outside of a species’ native range were identified using the Native Species Resolver (NSR; http://bien.nceas.ucsb.edu/bien/tools/nsr/), which uses published checklists and endemism data to determine if the observed species is native in a given location. Observations of potentially cultivated plants are flagged and removed from the observation data.

Additional steps were undertaken to prepare the cleaned data for species distribution modeling. To reduce spatial autocorrelation, only one record was retained if multiple records were found in a 5-km grid cell. Such thinning is recommended when modeling the occurrence density (in contrast to the density of individuals in which retaining each record is critical) (*87*). The occurrences were further adjusted to ensure that all retained records were at least 10 km apart, using the default thinning algorithm of “spthin” (v.0.2.0) (*88*).

The geographic distributions of plant species were modeled using twelve climate and eight soil variables (correlation coefficients < 0.7) at a 5 km resolution. The twelve bioclimatic variables were obtained from CHELSA (*89*), including BIO1 (Annual Mean Temperature), BIO12 (Annual Precipitation), BIO14 (Precipitation of the Driest Month), BIO15 (Precipitation Seasonality-Coefficient of Variation), BIO18 (Precipitation of the Warmest Quarter), BIO19 (Precipitation of the Coldest Quarter), BIO2 (Mean Diurnal Range), BIO8 (Mean Temperature of the Wettest Quarter), Frost Change Frequency, Growing Season Length, Growing Season Temperature, and Snow Water Equivalent. The eight SoilGrids variables (*90*) were Cation Exchange Capacity, Coarse Fragments Volume, clay content in the soil, total nitrogen content, Organic Carbon Stock, pH in water, the percentage of silt content in the soil, and Soil Organic Carbon.

Three modeling techniques were employed based on the number of species occurrences. For plant species with at least 15 spatially unique records, we used Poisson point process models (a generalized version of MaxEnt) fit with the R package glmnet (v.4.0-2) (*91*)(70). For species with 3–14 spatially unique records, we used a range-bagging algorithm (*92*), which uses an ensemble of statistically generated convex hulls in environmental space. For any species with 1– 3 spatially unique records, we considered the cells in which records were found (at 5 km resolution) to be its geographic range. The selection of the algorithms is based on the philosophy of using a more conservative approach when there is fewer data for model training.

After modelling, we selected 35,775 plant species with geographic ranges within the Amazon basin area. To further investigate the exposure of metal contaminants on species directly associated with aquatic environments, we also selected 276 tree species occurring in either Igapo or Varzea forests in Amazonia, i.e., forests susceptible to flooding pulses, from the list of species made available by (*36*).

All the species had their geographic ranges cropped to the extension of the Amazonia basin as proposed by (*93*), which includes areas of the Cerrado Biome and montane regions that drain into the Amazon River. This delimitation of Amazonia encompasses an area of 7,595,000 km^2^ and includes areas from Brazil, Bolivia, Peru, Ecuador, Colombia, Venezuela, Guyana, Suriname, and French Guiana. The final numbers of vertebrate species included in the analysis were 744 mammals, 1,212 amphibians, 661 reptiles, and 2,361 birds, 2,150 fish species. Plants included vascular species only and constituted 35,775 species.

### Mining contamination along waterways

The rasters of metal contamination were built as follows. Registered (formal) inactive and active mines locations in the Amazon Region were taken from the Water and Planetary Health Analytics global mines database (WAPHA 2023) and attenuation of mean sediment concentration downstream of four toxic elements (As, Cu, Pb, Zn) associated with the mined commodity, calculated using the methods given in (*19*) in their global assessment of the impact of metal mining on river systems. For the current study we used the HydroSheds (v1.1) 15 arc-second DEM (*94*) to construct a gridded river network. We identified those 500m cells downstream of mines with predicted concentrations of each metal greater than or equal to the Probable Effect Level (PEL) concentrations for freshwater sediment established by Canadian Environmental Quality Guidelines (*35*).

Artisanal (informal) mine locations were derived from mining scars mapped by Amazon Mining Watch (https://amazonminingwatch.org) using Sentinel-2 satellite imagery. We used their open-source 2018-23 cumulative product (Earth Genome, 2024; accessed August 03, 2024). Due to rapid vegetation regrowth, older mining scars from before 2018 may be undetected, so these data will likely underestimate contamination sources. Mining scar polygons were converted to 500m-spaced points, then snapped to the nearest 500 m grid cell of the river network to estimate the river reach exposed to artisanal mining. The mining scar product does not discriminate between artisanal gold mining and tin (cassiterite) mining, as both are found in alluvial placer deposits. The region’s registered mine locations showed a predominance of gold (*n*=937) and tin mines (*n*=340). Most mines were spatially aggregated in the same areas, thus likely affecting the same river reaches.

Consequently, without data allowing further discrimination, we treated all the river locations as sources of mercury contamination from gold mining. However, this may have led to local overestimation. The only published analysis of Hg attenuation from a mine downriver that we are aware of is presented by (*95*), but this is from sources with relatively low Hg concentrations (maximum 5.96 mg/kg) compared to those found in sediment/soil near artisanal gold mining using mercury as an amalgam [(*96*)], ∼70-90 mg/kg Hg) and very high values in tailing waste, which is directly discharged into waterways (*97*). Similar source Hg in sediment levels (57.1 mg/kg) arising from historical industrial contamination were found in a study by (*98*) along a waterway in Tennessee, who also measured the concentration of Hg in river sediment by distance from this source. We used these data to statistically model Hg concentration as the dependent variable against distance from source in a 2-parameter negative exponential regression: Hg = 29.0760 * *e* distance * −0.0446 with Hg measured in mg/kg and distance in km (R^2^ = 0.226, RMSE = 9.241). As for Registered Mines, we identified 500m cells downstream of artisanal gold mining mercury contamination sources with predicted Hg concentrations greater than or equal to the Probable Effect Level (PEL) concentrations for freshwater sediment established by Canadian Environmental Quality Guidelines (*35*).

Environmental hazards associated with metals in mining sediments are assessed using threshold values like the Threshold Effect Level (TEL) and Probable Effect Level (PEL), which help predict biological impacts. The TEL indicates the concentration above which metals may cause rare adverse effects, while the PEL represents the level at which clear adverse effects are expected. The (*35*) have set short- and long-term exposure thresholds for aquatic environments to signal unacceptable ecological impacts. While local studies in the Amazon have reported metal contamination exceeding these thresholds (*29*, *99*), regional assessments remain limited. We consider that the environmental hazards posed by metal contamination in the environment are considered toxic values above the PEL threshold. Distance downstream from mine location predicted to exceed or equal PEL concentrations with 95%CI were As (92.0 +/− 36.23 km), Pb (28.77 +/− 4.85 km), Cu (10.17 +/− 4.03 km), Zn (8.63 +/− 1.44 km) and Hg (91.74 [46.15 - 480.61] km).

Spatial data were projected to UTM zone 20S (EPSG:32720) at a grid cell length of 500m. Mine contamination was analyzed using MATLAB (*100*) and QGIS 3.34 Prizen (*101*).

### Calculating species ranges exposure to mining contamination

We calculated the fraction of the species geographic ranges that overlap with the metal contamination rasters of As, Cu, Hg, Pb, and Zn (Figure 1B). The higher the overlap with the metal contamination, the higher the exposure of each species to the metals. We focused on a single threshold of exposure in the main text. We considered species with >=1% of their range exposed to omit species with such little exposure that they would likely not be of concern. Based on a preliminary assessment of each species exposures, we do not emphasize our results in the thresholds of >=25%, >=50%, and >=75%, as previously recommended (*102*). This is because contrary to other threats such as increasing temperature or deforestation, contamination in aquatic environments has less spatial intersection with large ranges, and therefore very few species had exposures above 25%.

### Comparing species exposure to mining and to fire

We used the results of the study of (*40*) to compare the exposure of plant species to fire and to metal contamination. The assessment of the compounding exposed between metal contamination and fire exposure is only possible because the study by Feng et al (39) employs a similar analytical approach to ours, utilizing species geographic ranges from the same source (30), and therefore, is to some extent comparable to our findings.

### Mapping contamination on diversity hotspots and indigenous territories

Diversity hotspots were defined as species richness quantiles and calculated by extracting the pixels that had the top 50 %, 25%, and 10% of the species richness maps of each taxa (Figure S1). Those diversity hotspots regions were then overlapped with a mosaic of metal contamination rasters with all five metals to calculate the fraction of hotspots in contaminated areas. Because Amazonia can be regionalize in biogeographical regions (*103*), we also addressed the fraction of the diversity hotspots that overlap with regions contaminated in the biogeochemical provinces as defined by (*104*)): Northwestern Amazonia (NWA), Southwestern Amazonia (SWA), Southern Amazonia (SA), Central Amazonia (CA), Northern Amazonia (NoA), and Eastern Amazonia (EA).

The spatial polygons of Indigenous territories were downloaded from (*105*). We evaluated the exposure of indigenous territories by calculating the maximum pixel value of a range exposure map using all species. The range exposure map was constructed by assigning the corresponding values to their exposed areas (one value for each species geographic range) and then calculating the median values per each 5 km by 5 km grid cell.

All spatial analyses were done in R (*106*) using functions of the R package terra (*107*)

## References and Notes

1. P. Potapov, M. C. Hansen, L. Laestadius, S. Turubanova, A. Yaroshenko, C. Thies, W. Smith, I. Zhuravleva, A. Komarova, S. Minnemeyer, E. Esipova, The last frontiers of wilderness: Tracking loss of intact forest landscapes from 2000 to 2013. Sci. Adv. 3, e1600821 (2017).

2. J. R. Allan, O. Venter, J. E. M. Watson, Temporally inter-comparable maps of terrestrial wilderness and the Last of the Wild. Sci. Data 4, 170187 (2017).

3. D. Armenteras, E. Berenguer, C. S. Andreazzi, L. M. Dávalos, F. Duponchelle, S. Hacon, A. G. Lescano, M. Macedo, N. Nascimento, “Chapter 21: Human well-being and health impacts of the degradation of terrestrial and aquatic ecosystems” in Amazon Assessment Report 2021 (UN Sustainable Development Solutions Network (SDSN), 2021; 10.55161/kryi5458).

4. B. Flores, E. Montoya, B. Sakschewski, N. Nascimento, A. Staal, R. A. Betts, C. Levis, D. Lapola, A. Esquível-Muelbert, C. C. Jakovac, C. A. Nobre, R. S. Oliveira, L. Borma, D. Nian, N. Boers, S. B. Hecht, H. ter Steege, J. Arieira, I. L. Lucas, E. Berenguer, J. Marengo, L. V. Gatti, C. R. C. Mattos, M. Hirota, Critical transitions in the Amazon forest system. Nature 626, 555–564 (2024).

5. E. N. Dethier, M. Silman, J. D. Leiva, S. Alqahtani, L. E. Fernandez, P. Pauca, S. Çamalan, P. Tomhave, F. J. Magilligan, C. E. Renshaw, D. A. Lutz, A global rise in alluvial mining increases sediment load in tropical rivers. Nature 620, 787–793 (2023).

6. J. Siqueira-Gay, J. P. Metzger, L. E. Sánchez, L. J. Sonter, Strategic planning to mitigate mining impacts on protected areas in the Brazilian Amazon. Nat. Sustain. 5, 853–860 (2022).

7. IPBES, Global assessment report on biodiversity and ecosystem services of the Intergovernmental Science-Policy Platform on Biodiversity and Ecosystem Services. IPBES secretariat, Bonn, Germany [Preprint] (2019). 10.5281/ZENODO.6417333.

8. P. Fearnside, E. Berenguer, D. Armenteras, F. Duponchelle, F. Mosquera Guerra, C. N. Jenkins, P. Bynoe, R. García-Villacorta, M. Macedo, A. L. Val, V. M. Fonseca de Almeida-Val, N. Nascimento, “Chapter 20: Drivers and impacts of changes in aquatic ecosystems” in Amazon Assessment Report 2021 (UN Sustainable Development Solutions Network (SDSN), 2021; 10.55161/idmb5770).

9. European Commission, EU Biodiversity Strategy for 2030: Bringing Nature Back into Our Lives (Publications Office of the European Union, 2021; https://data.europa.eu/doi/10.2779/677548).

10. L. Posthuma, J. van Gils, M. C. Zijp, D. van de Meent, D. de Zwart, Species sensitivity distributions for use in environmental protection, assessment, and management of aquatic ecosystems for 12 386 chemicals: Species sensitivity distributions for 12,386 compounds. Environ. Toxicol. Chem. 38, 905–917 (2019).

11. M. de Almeida Ribeiro Carvalho, W. G. Botero, L. C. de Oliveira, Natural and anthropogenic sources of potentially toxic elements to aquatic environment: a systematic literature review. Environ. Sci. Pollut. Res. Int. 29, 51318–51338 (2022).

12. G. Sigmund, M. Ågerstrand, A. Antonelli, T. Backhaus, T. Brodin, M. L. Diamond, W. R. Erdelen, D. C. Evers, T. Hofmann, T. Hueffer, A. Lai, J. P. M. Torres, L. Mueller, A. L. Perrigo, M. C. Rillig, A. Schaeffer, M. Scheringer, K. Schirmer, A. Tlili, A. Soehl, R. Triebskorn, P. Vlahos, C. Vom Berg, Z. Wang, K. J. Groh, Addressing chemical pollution in biodiversity research. Glob. Chang. Biol. 29, 3240–3255 (2023).

13. D. Hou, X. Jia, L. Wang, S. P. McGrath, Y.-G. Zhu, Q. Hu, F.-J. Zhao, M. S. Bank, D. O’Connor, J. Nriagu, Global soil pollution by toxic metals threatens agriculture and human health. Science 388, 316–321 (2025).

14. M. A. da Costa, F. J. Rios, The gold mining industry in Brazil: A historical overview. Ore Geol. Rev. 148, 105005 (2022).

15. C. Hoorn, F. P. Wesselingh, H. ter Steege, M. A. Bermudez, A. Mora, J. Sevink, I. Sanmartín, A. Sanchez-Meseguer, C. L. Anderson, J. P. Figueiredo, C. Jaramillo, D. Riff, F. R. Negri, H. Hooghiemstra, J. Lundberg, T. Stadler, T. Särkinen, A. Antonelli, Amazonia through time: Andean uplift, climate change, landscape evolution, and biodiversity. Science 330, 927–931 (2010).

16. L. M. Diele-Viegas, E. J. de A. L. Pereira, C. F. D. Rocha, The new Brazilian gold rush: Is Amazonia at risk? For. Policy Econ. 119, 102270 (2020).

17. S. Villén-Pérez, L. Anaya-Valenzuela, D. Conrado da Cruz, P. M. Fearnside, Mining threatens isolated indigenous peoples in the Brazilian Amazon. Glob. Environ. Change 72, 102398 (2022).

18. V. M. Prasniewski, W. González-Daza, G. do V. Alvarenga, L. Santos-Silva, A. L. Teixido, T. J. Izzo, Economic, environmental and social threats of a mining exploration proposal on indigenous lands of Brazil. Acta Amazon. 54, e54fo23192 (2024).

19. M. G. Macklin, C. J. Thomas, A. Mudbhatkal, P. A. Brewer, K. A. Hudson-Edwards, J. Lewin, P. Scussolini, D. Eilander, A. Lechner, J. Owen, G. Bird, D. Kemp, K. R. Mangalaa, Impacts of metal mining on river systems: a global assessment. Science 381, 1345–1350 (2023).

20. J. R. Gerson, N. Szponar, A. A. Zambrano, B. Bergquist, E. Broadbent, C. T. Driscoll, G. Erkenswick, D. C. Evers, L. E. Fernandez, H. Hsu-Kim, G. Inga, K. N. Lansdale, M. J. Marchese, A. Martinez, C. Moore, W. K. Pan, R. P. Purizaca, V. Sánchez, M. Silman, E. A. Ury, C. Vega, M. Watsa, E. S. Bernhardt, Amazon forests capture high levels of atmospheric mercury pollution from artisanal gold mining. Nat. Commun. 13, 559 (2022).

21. L. F. Viana, F. Kummrow, C. A. L. Cardoso, N. A. de Lima, J. C. J. Solórzano, B. do A. Crispim, A. Barufatti, A. C. Florentino, High concentrations of metals in the waters from Araguari River lower section (Amazon biome): Relationship with land use and cover, ecotoxicological effects and risks to aquatic biota. Chemosphere 285, 131451 (2021).

22. T. D. Jardine, K. A. Kidd, N. O’ Driscoll, Food web analysis reveals effects of pH on mercury bioaccumulation at multiple trophic levels in streams. Aquat. Toxicol. 132–133, 46–52 (2013).

23. A. V. Waichman, G. S. de S. Nunes, R. de Oliveira, I. López-Heras, A. Rico, Human health risks associated to trace elements and metals in commercial fish from the Brazilian Amazon. J. Environ. Sci. (China) 148, 230–242 (2025).

24. J. J. Melendez-Perez, A. H. Fostier, J. A. Carvalho Jr, C. C. Windmöller, J. C. Santos, A. Carpi, Soil and biomass mercury emissions during a prescribed fire in the Amazonian rain forest. Atmos. Environ. (1994) 96, 415–422 (2014).

25. M. Saaristo, T. Brodin, S. Balshine, M. G. Bertram, B. W. Brooks, S. M. Ehlman, E. S. McCallum, A. Sih, J. Sundin, B. B. M. Wong, K. E. Arnold, Direct and indirect effects of chemical contaminants on the behaviour, ecology and evolution of wildlife. Proc. Biol. Sci. 285, 20181297 (2018).

26. M. Michelangeli, J. M. Martin, N. Pinter-Wollman, C. C. Ioannou, E. S. McCallum, M. G. Bertram, T. Brodin, Predicting the impacts of chemical pollutants on animal groups. Trends Ecol. Evol. 37, 789–802 (2022).

27. L. A. Martinelli, Jose R. Ferreira, B. R. Forsberg, R. L. Victoria, Mercury contamination in the Amazon: A gold rush consequence. Ambio 17, 252–254 (1988).

28. S. E. Hook, E. P. Gallagher, G. E. Batley, The role of biomarkers in the assessment of aquatic ecosystem health: Biomarkers in the Assessment of Aquatic Ecosystem Health. Integr. Environ. Assess. Manag. 10, 327–341 (2014).

29. M. V. Capparelli, G. M. Moulatlet, D. M. de S. Abessa, O. Lucas-Solis, B. Rosero, E. Galarza, D. Tuba, N. Carpintero, V. Ochoa-Herrera, I. Cipriani-Avila, An integrative approach to identify the impacts of multiple metal contamination sources on the Eastern Andean foothills of the Ecuadorian Amazonia. Sci. Total Environ. 709, 136088 (2020).

30. R. P. Anderson, A. Townsend Peterson, E. Martinez-Meyer, J. Soberon, M. Nakamura, R. G. Pearson, M. B. Araujo, Ecological Niches and Geographic Distributions (MPB-49) (Princeton University Press, Princeton, NJ, 2017; https://www.jstor.org/stable/j.ctt7stnh) Monographs in Population Biology.

31. M. G. Macklin, P. A. Brewer, K. A. Hudson-Edwards, G. Bird, T. J. Coulthard, I. A. Dennis, P. J. Lechler, J. R. Miller, J. N. Turner, A geomorphological approach to the management of rivers contaminated by metal mining. Geomorphology (Amst.) 79, 423–447 (2006).

32. V. J. Isaac, M. C. Almeida, T. Giarrizzo, C. P. Deus, R. Vale, G. Klein, A. Begossi, Food consumption as an indicator of the conservation of natural resources in riverine communities of the Brazilian Amazon. An. Acad. Bras. Cienc. 87, 2229–2242 (2015).

33. B. S. Maitner, B. Boyle, N. Casler, R. Condit, J. Donoghue, S. M. Durán, D. Guaderrama, C. E. Hinchliff, P. M. Jørgensen, N. J. B. Kraft, B. McGill, C. Merow, N. Morueta Holme, R. K. Peet, B. Sandel, M. Schildhauer, S. A. Smith, J. Svenning, B. Thiers, C. Violle, S. Wiser, B. J. Enquist, The bien r package: A tool to access the Botanical Information and Ecology Network (BIEN) database. Methods in Ecology and Evolution 9, 373–379 (2018).

34. M. Lumbierres, P. R. Dahal, C. D. Soria, M. Di Marco, S. H. M. Butchart, P. F. Donald, C. Rondinini, Area of Habitat maps for the world’s terrestrial birds and mammals. Sci. Data 9, 749 (2022).

35. CCME, Sediment Quality Guidelines for the Protection of Aquatic Life Freshwater and (2024).

36. J. E. Householder, F. Wittmann, J. Schöngart, M. T. F. Piedade, W. J. Junk, E. M. Latrubesse, A. C. Quaresma, L. O. Demarchi, G. de S Lobo, D. P. P. de Aguiar, R. L. Assis, A. Lopes, P. Parolin, I. Leão do Amaral, L. de S. Coelho, F. D. de Almeida Matos, D. de A. Lima Filho, R. P. Salomão, C. V. Castilho, J. E. Guevara-Andino, M. de J. V. Carim, O. L. Phillips, D. Cárdenas López, W. E. Magnusson, D. Sabatier, J. D. C. Revilla, J.-F. Molino, M. V. Irume, M. P. Martins, J. R. da S. Guimarães, J. F. Ramos, D. de J. Rodrigues, O. S. Bánki, C. A. Peres, N. C. A. Pitman, J. E. Hawes, E. J. Almeida, L. F. Barbosa, L. Cavalheiro, M. C. V. Dos Santos, B. G. Luize, E. M. M. de L. Novo, P. Núñez Vargas, T. S. F. Silva, E. M. Venticinque, A. G. Manzatto, N. F. C. Reis, J. Terborgh, K. R. Casula, F. R. C. Costa, E. N. Honorio Coronado, A. Monteagudo Mendoza, J. C. Montero, T. R. Feldpausch, G. A. Aymard C, C. Baraloto, N. Castaño Arboleda, J. Engel, P. Petronelli, C. E. Zartman, T. J. Killeen, L. M. Rincón, B. S. Marimon, B. H. Marimon-Junior, J. Schietti, T. R. Sousa, R. Vasquez, B. Mostacedo, D. Dantas do Amaral, H. Castellanos, M. B. de Medeiros, M. F. Simon, A. Andrade, J. L. Camargo, W. F. Laurance, S. G. W. Laurance, E. de S. Farias, M. A. Lopes, J. L. L. Magalhães, H. E. Mendonça Nascimento, H. L. de Queiroz, R. Brienen, P. R. Stevenson, A. Araujo-Murakami, T. R. Baker, B. B. L. Cintra, Y. O. Feitosa, H. F. Mogollón, J. C. Noronha, F. R. Barbosa, R. de Sá Carpanedo, J. F. Duivenvoorden, M. R. Silman, L. V. Ferreira, C. Levis, J. R. Lozada, J. A. Comiskey, F. C. Draper, J. J. de Toledo, G. Damasco, N. Dávila, R. García-Villacorta, A. Vicentini, F. Cornejo Valverde, A. Alonso, L. Arroyo, F. Dallmeier, V. H. F. Gomes, E. M. Jimenez, D. Neill, M. C. Peñuela Mora, F. A. Carvalho, F. Coelho de Souza, K. J. Feeley, R. Gribel, M. P. Pansonato, M. Ríos Paredes, J. Barlow, E. Berenguer, K. G. Dexter, J. Ferreira, P. V. A. Fine, M. C. Guedes, I. Huamantupa-Chuquimaco, J. C. Licona, T. Pennington, B. E. Villa Zegarra, V. A. Vos, C. Cerón, É. Fonty, T. W. Henkel, P. Maas, E. Pos, M. Silveira, J. Stropp, R. Thomas, D. Daly, W. Milliken, G. Pardo Molina, I. C. G. Vieira, B. W. Albuquerque, W. Campelo, T. Emilio, A. Fuentes, B. Klitgaard, J. L. Marcelo Pena, P. F. Souza, J. S. Tello, C. Vriesendorp, J. Chave, A. Di Fiore, R. R. Hilário, L. de O. Pereira, J. F. Phillips, G. Rivas-Torres, T. R. van Andel, P. von Hildebrand, W. Balee, E. M. Barbosa, L. C. de M. Bonates, H. P. D. Doza, R. Z. Gómez, T. Gonzales, G. P. G. Gonzales, B. Hoffman, A. B. Junqueira, Y. Malhi, I. P. de A. Miranda, L. F. Mozombite-Pinto, A. Prieto, A. Rudas, A. R. Ruschel, N. Silva, C. I. A. Vela, S. Zent, E. L. Zent, A. Cano, Y. A. Carrero Márquez, D. F. Correa, J. B. P. Costa, B. M. Flores, D. Galbraith, M. Holmgren, M. Kalamandeen, M. T. Nascimento, A. A. Oliveira, H. Ramirez-Angulo, M. Rocha, V. V. Scudeller, R. Sierra, M. Tirado, M. N. Umaña, G. van der Heijden, E. Vilanova Torre, M. A. Ahuite Reategui, C. Baider, H. Balslev, S. Cárdenas, L. F. Casas, W. Farfan-Rios, C. Ferreira, R. Linares-Palomino, C. Mendoza, I. Mesones, G. A. Parada, A. Torres-Lezama, L. E. Urrego Giraldo, D. Villarroel, R. Zagt, M. N. Alexiades, E. A. de Oliveira, K. Garcia-Cabrera, L. Hernandez, W. Palacios Cuenca, S. Pansini, D. Pauletto, F. Ramirez Arevalo, A. F. Sampaio, E. H. Valderrama Sandoval, L. Valenzuela Gamarra, H. Ter Steege, One sixth of Amazonian tree diversity is dependent on river floodplains. Nat. Ecol. Evol. 8, 901–911 (2024).

37. C. D. Foy, R. T. Chaney, M. C. White, The physiology of metal toxicity in plants. Annual review of plant physiology 29, 511–566 (1978).

38. G. P. Cobb, K. Sands, M. Waters, B. G. Wixson, E. Dorward-King, Accumulation of heavy metals by vegetables grown in mine wastes. Environ. Toxicol. Chem. 19, 600–607 (2000).

39. F. Wittmann, A. de O. Wittmann, “Use of amazonian floodplain trees” in Ecological Studies (Springer Netherlands, Dordrecht, 2010; 10.1007/978-90-481-8725-6_19)Ecological studies: analysis and synthesis. Berlin, Heidelberg, New York NY, pp. 389–418.

40. X. Feng, C. Merow, Z. Liu, D. S. Park, P. Roehrdanz, B. S. Maitner, E. A. Newman, B. Boyle, A. M. Lien, J. R. Burger, M. Pires, P. Brando, M. Bush, C. McMichael, D. M. Neves, E. Nikolopoulos, S. Saleska, L. Hannah, D. Breshears, T. Evans, J. R. Soto, K. Ernst, B. Enquist, How deregulation, drought and increasing fire impact Amazonian biodiversity. Nature 597, 516–521 (2021).

41. H. Ter Steege, N. C. A. Pitman, T. J. Killeen, W. F. Laurance, C. A. Peres, J. E. Guevara, R. P. Salomão, C. V. Castilho, I. L. Amaral, F. D. de Almeida Matos, L. de Souza Coelho, W. E. Magnusson, O. L. Phillips, D. de Andrade Lima Filho, M. de Jesus Veiga Carim, M. V. Irume, M. P. Martins, J.-F. Molino, D. Sabatier, F. Wittmann, D. C. López, J. R. da Silva Guimarães, A. M. Mendoza, P. N. Vargas, A. G. Manzatto, N. F. C. Reis, J. Terborgh, K. R. Casula, J. C. Montero, T. R. Feldpausch, E. N. Honorio Coronado, A. J. D. Montoya, C. E. Zartman, B. Mostacedo, R. Vasquez, R. L. Assis, M. B. Medeiros, M. F. Simon, A. Andrade, J. L. Camargo, S. G. W. Laurance, H. E. M. Nascimento, B. S. Marimon, B.-H. Marimon Jr, F. Costa, N. Targhetta, I. C. G. Vieira, R. Brienen, H. Castellanos, J. F. Duivenvoorden, H. F. Mogollón, M. T. F. Piedade, G. A. Aymard C, J. A. Comiskey, G. Damasco, N. Dávila, R. García-Villacorta, P. R. S. Diaz, A. Vincentini, T. Emilio, C. Levis, J. Schietti, P. Souza, A. Alonso, F. Dallmeier, L. V. Ferreira, D. Neill, A. Araujo-Murakami, L. Arroyo, F. A. Carvalho, F. C. Souza, D. D. do Amaral, R. Gribel, B. G. Luize, M. P. Pansonato, E. Venticinque, P. Fine, M. Toledo, C. Baraloto, C. Cerón, J. Engel, T. W. Henkel, E. M. Jimenez, P. Maas, M. C. P. Mora, P. Petronelli, J. D. C. Revilla, M. Silveira, J. Stropp, R. Thomas-Caesar, T. R. Baker, D. Daly, M. R. Paredes, N. F. da Silva, A. Fuentes, P. M. Jørgensen, J. Schöngart, M. R. Silman, N. C. Arboleda, B. B. L. Cintra, F. C. Valverde, A. Di Fiore, J. F. Phillips, T. R. van Andel, P. von Hildebrand, E. M. Barbosa, L. C. de Matos Bonates, D. de Castro, E. de Sousa Farias, T. Gonzales, J.-L. Guillaumet, B. Hoffman, Y. Malhi, I. P. de Andrade Miranda, A. Prieto, A. Rudas, A. R. Ruschell, N. Silva, C. I. A. Vela, V. A. Vos, E. L. Zent, S. Zent, A. Cano, M. T. Nascimento, A. A. Oliveira, H. Ramirez-Angulo, J. F. Ramos, R. Sierra, M. Tirado, M. N. U. Medina, G. van der Heijden, E. V. Torre, C. Vriesendorp, O. Wang, K. R. Young, C. Baider, H. Balslev, N. de Castro, W. Farfan-Rios, C. Ferreira, C. Mendoza, I. Mesones, A. Torres-Lezama, L. E. U. Giraldo, D. Villarroel, R. Zagt, M. N. Alexiades, K. Garcia-Cabrera, L. Hernandez, I. Huamantupa-Chuquimaco, W. Milliken, W. P. Cuenca, S. Pansini, D. Pauletto, F. R. Arevalo, A. F. Sampaio, E. H. Valderrama Sandoval, L. V. Gamarra, Estimating the global conservation status of more than 15,000 Amazonian tree species. Sci. Adv. 1, e1500936 (2015).

42. W. J. Junk, S. M. G. M., P. B. Bayley, Freshwater fishes of the Amazon River basin: their biodiversity, fisheries, and habitats. Aquatic Ecosystem Health & Management 10, 153–173 (2007).

43. P. C. Basta, A. C. D. de Vasconcellos, G. Hallwass, D. Yokota, D. de Oliveira d’El Rei Pinto, D. S. de Aguiar, C. C. de Souza, M. Oliveira-da-Costa, Risk assessment of mercury-contaminated fish consumption in the Brazilian Amazon: An ecological study. Toxics 11 (2023).

44. L. Córdoba-Tovar, J. Marrugo-Negrete, P. R. Barón, S. Díez, Drivers of biomagnification of Hg, As and Se in aquatic food webs: A review. Environ. Res. 204, 112226 (2022).

45. P. Wu, M. J. Kainz, A. G. Bravo, S. Åkerblom, L. Sonesten, K. Bishop, The importance of bioconcentration into the pelagic food web base for methylmercury biomagnification: A meta-analysis. Sci. Total Environ. 646, 357–367 (2019).

46. M. E. Crespo-Lopez, M. Augusto-Oliveira, A. Lopes-Araújo, L. Santos-Sacramento, P. Yuki Takeda, B. de M. Macchi, J. L. M. do Nascimento, C. S. F. Maia, R. R. Lima, G. P. Arrifano, Mercury: What can we learn from the Amazon? Environ. Int. 146, 106223 (2021).

47. C. A. Sayer, E. Fernando, R. R. Jimenez, N. B. W. Macfarlane, G. Rapacciuolo, M. Böhm, T. M. Brooks, T. Contreras-MacBeath, N. A. Cox, I. Harrison, M. Hoffmann, R. Jenkins, K. G. Smith, J.-C. Vié, J. C. Abbott, D. J. Allen, G. R. Allen, V. Barrios, J.-P. Boudot, S. F. Carrizo, P. Charvet, V. Clausnitzer, L. Congiu, K. A. Crandall, N. Cumberlidge, A. Cuttelod, J. Dalton, A. G. Daniels, S. De Grave, G. De Knijf, K.-D. B. Dijkstra, R. A. Dow, J. Freyhof, N. García, J. Gessner, A. Getahun, C. Gibson, M. J. Gollock, M. I. Grant, A. E. R. Groom, M. P. Hammer, G. A. Hammerson, C. Hilton-Taylor, L. Hodgkinson, R. A. Holland, R. W. Jabado, D. Juffe Bignoli, V. J. Kalkman, B. K. Karimov, J. Kipping, M. Kottelat, P. A. Lalèyè, H. K. Larson, M. Lintermans, F. Lozano, A. Ludwig, T. J. Lyons, L. Máiz-Tomé, S. Molur, H. H. Ng, C. Numa, A. F. Palmer-Newton, C. Pike, H. E. Pippard, C. N. M. Polaz, C. M. Pollock, R. Raghavan, P. S. Rand, T. Ravelomanana, R. E. Reis, C. L. Rigby, J. A. Scott, P. H. Skelton, M. R. Sloat, J. Snoeks, M. L. J. Stiassny, H. H. Tan, Y. Taniguchi, E. B. Thorstad, M. F. Tognelli, A. G. Torres, Y. Torres, D. Tweddle, K. Watanabe, J. R. S. Westrip, E. G. E. Wright, E. Zhang, W. R. T. Darwall, One-quarter of freshwater fauna threatened with extinction. Nature 638, 138–145 (2025).

48. G. M. Moulatlet, N. Yacelga, A. Rico, A. Mora, R. A. Hauser-Davis, M. Cabrera, M. V. Capparelli, A systematic review on metal contamination due to mining activities in the Amazon basin and associated environmental hazards. Chemosphere 339, 139700 (2023).

49. G. Azambuja, I. Kaefer, A. L. Val, D. Kochhann, “A double threat to amazonian amphibian larvae: A review of toxic contaminants and their interaction with global climate change” in The Future of Amazonian Aquatic Biota (Springer Nature Switzerland, Cham, 2024; 10.1007/978-3-031-66822-7_9), pp. 271–311.

50. S. E. Diringer, A. J. Berky, M. Marani, E. J. Ortiz, O. Karatum, D. L. Plata, W. K. Pan, H. Hsu-Kim, Deforestation due to artisanal and small-scale gold mining exacerbates soil and mercury mobilization in Madre de Dios, Peru. Environ. Sci. Technol. 54, 286–296 (2020).

51. C. Levis, B. M. Flores, J. V. Campos-Silva, N. Peroni, A. Staal, M. C. G. Padgurschi, W. Dorshow, B. Moraes, M. Schmidt, T. W. Kuikuro, H. Kuikuro, K. Wauja, K. Kuikuro, A. Kuikuro, C. Fausto, B. Franchetto, J. Watling, H. Lima, M. Heckenberger, C. R. Clement, Contributions of human cultures to biodiversity and ecosystem conservation. Nat. Ecol. Evol. 8, 866–879 (2024).

52. L. Wyatt, E. J. Ortiz, B. Feingold, A. Berky, S. Diringer, A. M. Morales, E. R. Jurado, H. Hsu-Kim, W. Pan, Spatial, temporal, and dietary variables associated with elevated mercury exposure in Peruvian riverine communities upstream and downstream of artisanal and small-scale gold mining. Int. J. Environ. Res. Public Health 14, 1582 (2017).

53. J. N. Pisconte, C. M. Vega, C. J. Sayers 2nd, C. S. Sevillano-Ríos, M. Pillaca, E. Quispe, V. Tejeda, C. Ascorra, M. R. Silman, L. E. Fernandez, Elevated mercury exposure in bird communities inhabiting Artisanal and Small-Scale Gold Mining landscapes of the southeastern Peruvian Amazon. Ecotoxicology 33, 472–483 (2024).

54. J. R. Peralta-Videa, M. L. Lopez, M. Narayan, G. Saupe, J. Gardea-Torresdey, The biochemistry of environmental heavy metal uptake by plants: implications for the food chain. Int. J. Biochem. Cell Biol. 41, 1665–1677 (2009).

55. W. Mertz, The essential trace elements. Science 213, 1332–1338 (1981).

56. H. Ali, E. Khan, M. A. Sajad, Phytoremediation of heavy metals--concepts and applications. Chemosphere 91, 869–881 (2013).

57. M. N. Rana, J. Tangpong, M. M. Rahman, Toxicodynamics of Lead, Cadmium, Mercury and Arsenic-induced kidney toxicity and treatment strategy: A mini review. Toxicol. Rep. 5, 704–713 (2018).

58. Z. Rahman, V. P. Singh, The relative impact of toxic heavy metals (THMs) (arsenic (As), cadmium (Cd), chromium (Cr)(VI), mercury (Hg), and lead (Pb)) on the total environment: an overview. Environ. Monit. Assess. 191, 419 (2019).

59. L. D. Lacerda, M. de Souza, M. G. Ribeiro, The effects of land use change on mercury distribution in soils of Alta Floresta, Southern Amazon. Environ. Pollut. 129, 247–255 (2004).

60. N. Szponar, C. M. Vega, J. Gerson, D. S. McLagan, M. Pillaca, S. Delgado, D. Lee, N. Rahman, L. E. Fernandez, E. S. Bernhardt, A. M. Kiefer, C. P. J. Mitchell, F. Wania, B. A. Bergquist, Tracing atmospheric mercury from artisanal and small-scale gold mining. Environ. Sci. Technol. 59, 5021–5033 (2025).

61. A. K. Silman, R. Chhabria, G. W. Hafzalla, L. Giffin, K. Kucharski, K. Myers, C. Culquichicón, S. Montero, A. G. Lescano, C. M. Vega, L. E. Fernandez, M. R. Silman, M. J. Kane, J. W. Sanders, Impairment in working memory and executive function associated with mercury exposure in indigenous populations in upper amazonian Peru. Int. J. Environ. Res. Public Health 19, 10989 (2022).

62. R. Adler Miserendino, J. R. D. Guimarães, G. Schudel, S. Ghosh, J. M. Godoy, E. K. Silbergeld, P. S. J. Lees, B. A. Bergquist, Mercury pollution in Amapá, Brazil: Mercury amalgamation in artisanal and small-scale gold mining or land-cover and land-use changes? ACS Earth Space Chem. 2, 441–450 (2018).

63. E. Galarza, G. M. Moulatlet, A. Rico, M. Cabrera, V. Pinos-Velez, A. Pérez-González, M. V. Capparelli, Human health risk assessment of metals and metalloids in mining areas of the Northeast Andean foothills of the Ecuadorian Amazon. Integr. Environ. Assess. Manag. 19, 706–716 (2023).

64. T. Toivonen, S. Mäki, R. Kalliola, The riverscape of Western Amazonia – a quantitative approach to the fluvial biogeography of the region: The riverscape of Western Amazonia. J. Biogeogr. 34, 1374–1387 (2007).

65. K. Ruokolainen, G. M. Moulatlet, G. Zuquim, C. Hoorn, H. Tuomisto, Geologically recent rearrangements in central Amazonian river network and their importance for the riverine barrier hypothesis. Front. Biogeogr. 11 (2019).

66. S. Obiri, P. O. Yeboah, S. Osae, S. Adu-Kumi, Levels of arsenic, mercury, cadmium, copper, lead, zinc and manganese in serum and whole blood of resident adults from mining and non-mining communities in Ghana. Environ. Sci. Pollut. Res. Int. 23, 16589–16597 (2016).

67. A. Begossi, S. V. Salivonchyk, G. Hallwass, N. Hanazaki, P. F. M. Lopes, R. A. M. Silvano, D. Dumaresq, J. Pittock, Fish consumption on the Amazon: a review of biodiversity, hydropower and food security issues. Braz. J. Biol. 79, 345–357 (2019).

68. S. A. Heilpern, R. DeFries, K. Fiorella, A. Flecker, S. A. Sethi, M. Uriarte, S. Naeem, Declining diversity of wild-caught species puts dietary nutrient supplies at risk. Sci. Adv. 7, eabf9967 (2021).

69. M. Cabrera, M. V. Capparelli, C. Ñacato-Ch, G. M. Moulatlet, I. López-Heras, M. Díaz González, D. Alvear-S, A. Rico, Effects of intensive agriculture and urbanization on water quality and pesticide risks in freshwater ecosystems of the Ecuadorian Amazon. Chemosphere 337, 139286 (2023).

70. J. L. Wilkinson, A. B. A. Boxall, D. W. Kolpin, K. M. Y. Leung, R. W. S. Lai, C. Galbán-Malagón, A. D. Adell, J. Mondon, M. Metian, R. A. Marchant, A. Bouzas-Monroy, A. Cuni-Sanchez, A. Coors, P. Carriquiriborde, M. Rojo, C. Gordon, M. Cara, M. Moermond, T. Luarte, V. Petrosyan, Y. Perikhanyan, C. S. Mahon, C. J. McGurk, T. Hofmann, T. Kormoker, V. Iniguez, J. Guzman-Otazo, J. L. Tavares, F. Gildasio De Figueiredo, M. T. P. Razzolini, V. Dougnon, G. Gbaguidi, O. Traoré, J. M. Blais, L. E. Kimpe, M. Wong, D. Wong, R. Ntchantcho, J. Pizarro, G.-G. Ying, C.-E. Chen, M. Páez, J. Martínez-Lara, J.-P. Otamonga, J. Poté, S. A. Ifo, P. Wilson, S. Echeverría-Sáenz, N. Udikovic-Kolic, M. Milakovic, D. Fatta-Kassinos, L. Ioannou-Ttofa, V. Belušová, J. Vymazal, M. Cárdenas-Bustamante, B. A. Kassa, J. Garric, A. Chaumot, P. Gibba, I. Kunchulia, S. Seidensticker, G. Lyberatos, H. P. Halldórsson, M. Melling, T. Shashidhar, M. Lamba, A. Nastiti, A. Supriatin, N. Pourang, A. Abedini, O. Abdullah, S. S. Gharbia, F. Pilla, B. Chefetz, T. Topaz, K. M. Yao, B. Aubakirova, R. Beisenova, L. Olaka, J. K. Mulu, P. Chatanga, V. Ntuli, N. T. Blama, S. Sherif, A. Z. Aris, L. J. Looi, M. Niang, S. T. Traore, R. Oldenkamp, O. Ogunbanwo, M. Ashfaq, M. Iqbal, Z. Abdeen, A. O’Dea, J. M. Morales-Saldaña, M. Custodio, H. de la Cruz, I. Navarrete, F. Carvalho, A. B. Gogra, B. M. Koroma, V. Cerkvenik-Flajs, M. Gombač, M. Thwala, K. Choi, H. Kang, J. L. C. Ladu, A. Rico, P. Amerasinghe, A. Sobek, G. Horlitz, A. K. Zenker, A. C. King, J.-J. Jiang, R. Kariuki, M. Tumbo, U. Tezel, T. T. Onay, J. B. Lejju, Y. Vystavna, Y. Vergeles, H. Heinzen, A. Pérez-Parada, D. B. Sims, M. Figy, D. Good, C. Teta, Pharmaceutical pollution of the world’s rivers. Proc. Natl. Acad. Sci. U. S. A. 119, e2113947119 (2022).

71. K. F. de Moraes, M. P. D. Santos, G. S. R. Gonçalves, G. L. de Oliveira, L. B. Gomes, M. G. M. Lima, Climate change and bird extinctions in the Amazon. PLoS One 15, e0236103 (2020).

72. K. J. Feeley, E. M. Rehm, B. Machovina, perspective: The responses of tropical forest species to global climate change: acclimate, adapt, migrate, or go extinct? Front. Biogeogr. 4 (2012).

73. G. P. Asner, W. Llactayo, R. Tupayachi, E. R. Luna, Elevated rates of gold mining in the Amazon revealed through high-resolution monitoring. Proc. Natl. Acad. Sci. U. S. A. 110, 18454–18459 (2013).

74. E. N. Dethier, M. R. Silman, L. E. Fernandez, J. C. Espejo, S. Alqahtani, P. Pauca, D. A. Lutz, Operation mercury: Impacts of national level armed forces intervention and anticorruption strategy on artisanal gold mining and water quality in the Peruvian Amazon. Conserv. Lett. 16 (2023).

75. D. Abessa, A. Famá, L. Buruaem, The systematic dismantling of Brazilian environmental laws risks losses on all fronts. Nat. Ecol. Evol. 3, 510–511 (2019).

76. P. K. Maurya, D. S. Malik, K. K. Yadav, A. Kumar, S. Kumar, H. Kamyab, Bioaccumulation and potential sources of heavy metal contamination in fish species in River Ganga basin: Possible human health risks evaluation. Toxicol. Rep. 6, 472–481 (2019).

77. L. Cortinhas Ferreira Neto, C. G. Diniz, R. V. Maretto, C. Persello, M. L. Silva Pinheiro, M. C. Castro, L. W. R. Sadeck, A. F. Filho, J. Cansado, A. A. de A. Souza, J. P. Feitosa, D. C. Santos, M. Adami, P. W. M. Souza-Filho, A. Stein, A. Biehl, A. Klautau, Uncontrolled illegal mining and garimpo in the Brazilian Amazon. Nat. Commun. 15, 9847 (2024).

78. A. B. García-Andrade, J. D. Carvajal-Quintero, P. A. Tedesco, F. Villalobos, Evolutionary and environmental drivers of species richness in poeciliid fishes across the Americas. Glob. Ecol. Biogeogr. 30, 1245–1257 (2021).

79. P. A. Tedesco, O. Beauchard, R. Bigorne, S. Blanchet, L. Buisson, L. Conti, J.-F. Cornu, M. S. Dias, G. Grenouillet, B. Hugueny, C. Jézéquel, F. Leprieur, S. Brosse, T. Oberdorff, A global database on freshwater fish species occurrence in drainage basins. Sci. Data 4, 170141 (2017).

80. B. Lehner, G. Grill, Global river hydrography and network routing: baseline data and new approaches to study the world’s large river systems: GLOBAL RIVER HYDROGRAPHY AND NETWORK ROUTING. Hydrol. Process. 27, 2171–2186 (2013).

81. M. Lumbierres, P. R. Dahal, M. Di Marco, S. H. M. Butchart, P. F. Donald, C. Rondinini, Translating habitat class to land cover to map area of habitat of terrestrial vertebrates. Conserv. Biol. 36, e13851 (2022).

82. R. Bivand, S. Pawley, rgrass: Interface Between “GRASS” Geographical Information System and “R.” [Preprint] (2025). https://github.com/osgeo/rgrass.

83. M. Buchhorn, B. Smets, L. Bertels, B. De Roo, M. Lesiv, N.-E. Tsendbazar, M. Herold, S. Fritz, Copernicus global land service: land cover 100m: collection 3: epoch 2015: globe. Version V3. 0. 1)[Data set] (2020).

84. T. G. Farr, P. A. Rosen, E. Caro, R. Crippen, The shuttle radar topography mission. doi: 10.1029/2005RG000183 (2007).

85. B. Boyle, N. Hopkins, Z. Lu, J. A. Raygoza Garay, D. Mozzherin, T. Rees, N. Matasci, M. L. Narro, W. H. Piel, S. J. McKay, S. Lowry, C. Freeland, R. K. Peet, B. J. Enquist, The taxonomic name resolution service: an online tool for automated standardization of plant names. BMC Bioinformatics 14, 16 (2013).

86. B. L. Boyle, B. S. Maitner, G. G. C. Barbosa, R. K. Sajja, X. Feng, C. Merow, E. A. Newman, D. S. Park, P. R. Roehrdanz, B. J. Enquist, Geographic name resolution service: A tool for the standardization and indexing of world political division names, with applications to species distribution modeling. PLoS One 17, e0268162 (2022).

87. S. J. Phillips, M. Dudík, Modeling of species distributions with Maxent: new extensions and a comprehensive evaluation. Ecography (Cop.) 31, 161–175 (2008).

88. M. E. Aiello-Lammens, R. A. Boria, A. Radosavljevic, B. Vilela, R. P. Anderson, spThin: an R package for spatial thinning of species occurrence records for use in ecological niche models. Ecography (Cop.) 38, 541–545 (2015).

89. D. N. Karger, M. P. Nobis, S. Normand, C. H. Graham, N. E. Zimmermann, CHELSA-TraCE21k–high-resolution (1 km) downscaled transient temperature and precipitation data since the Last Glacial Maximum. Climate of the Past 19, 439–456 (2023).

90. T. Hengl, J. Mendes de Jesus, G. B. M. Heuvelink, M. Ruiperez Gonzalez, M. Kilibarda, A. Blagotić, W. Shangguan, M. N. Wright, X. Geng, B. Bauer-Marschallinger, M. A. Guevara, R. Vargas, R. A. MacMillan, N. H. Batjes, J. G. B. Leenaars, E. Ribeiro, I. Wheeler, S. Mantel, B. Kempen, SoilGrids250m: Global gridded soil information based on machine learning. PLoS One 12, e0169748 (2017).

91. J. Friedman, T. Hastie, R. Tibshirani, Regularization paths for generalized linear models via coordinate descent. J. Stat. Softw. 33, 1–22 (2010).

92. J. M. Drake, Range bagging: a new method for ecological niche modelling from presence-only data. J. R. Soc. Interface 12, 20150086 (2015).

93. H. D. Eva, O. Huber, F. Achard, H. Balslev, S. Beck, H. Behling, A. S. Belward, R. Beuchle, A. M. Cleef, M. Colchester, J. Duivenvoorden, M. Hoogmoed, W. Junk, P. Kabat, B. Kruijt, Y. Malhi, J. M. Müller, J. M. Pereira, C. Peres, G. T. Prance, J. Roberts, J. Salo, A Proposal for Defining the Geographical Boundaries of Amazonia; Synthesis of the Results from an Expert Consultation Workshop Organized by the European Commission in Collaboration with the Amazon Cooperation Treaty Organization - JRC Ispra, 7-8 June 2005 (European Commission, 2005)EUR.

94. B. Lehner, K. Verdin, A. Jarvis, New global hydrography derived from spaceborne elevation data. Eos (Washington DC) 89, 93–94 (2008).

95. L. Wei, M. Cai, Y. Du, J. Tang, Q. Wu, T. Xiao, Chen, Spatial attenuation of mining/smelting-derived metal pollution in sediments from tributaries of the Upper Han River, China. Mine Water and the Environment 38, 410–420 (2019).

96. P. S. Soe, W. T. Kyaw, K. Arizono, Y. Ishibashi, T. Agusa, Mercury pollution from artisanal and small-scale gold mining in Myanmar and other Southeast Asian countries. Int. J. Environ. Res. Public Health 19, 6290 (2022).

97. A. Yuliyanti, A. Aminuddin, “Mercury contamination in artisanal gold mining sites in Indonesia and the remediation” in Proceedings of the 3rd Sriwijaya International Conference on Environmental Issues, SRICOENV 2022, October 5th, 2022, Palembang, South Sumatera, Indonesia (EAI, 2023; 10.4108/eai.5-10-2022.2328332).

98. S. Brooks, V. Eller, J. Dickson, J. Earles, K. Lowe, T. Mehlhorn, T. Olsen, C. Derolph, D. Watson, D. Phillips, M. Peterson, Mercury Content of Sediments in East Fork Poplar Creek: Current Assessment and Past Trends (2017).

99. K. C. Carrillo, A. Rodríguez-Romero, A. Tovar-Sánchez, G. Ruiz-Gutiérrez, J. R. V. Fuente, Geochemical baseline establishment, contamination level and ecological risk assessment of metals and As in the Limoncocha lagoon sediments, Ecuadorian Amazon region. J. Soils Sediments 22, 293–315 (2022).

100. The MathWorks Inc, MATLAB and the MathWorks, Inc (American Institute of Aeronautics and Astronautics, Reston, VA, 2022; 10.2514/5.9781600861628.0425.0428).

101. QGIS Development Team, QGIS Geographic Information System (Open Source Geospatial Foundation Project, 2025; http://qgis.osgeo.org).

102. E. I. Ameca y Juárez, G. M. Mace, G. Cowlishaw, W. A. Cornforth, N. Pettorelli, Assessing exposure to extreme climatic events for terrestrial mammals. Conservation Letters 6, 145–153 (2013).

103. C. Dambros, G. Zuquim, G. M. Moulatlet, F. R. C. Costa, H. Tuomisto, C. C. Ribas, R. Azevedo, F. Baccaro, P. E. D. Bobrowiec, M. S. Dias, T. Emilio, H. M. V. Espirito-Santo, F. O. G. Figueiredo, E. Franklin, C. Freitas, M. B. Graça, F. d’Horta, R. P. Leitão, M. Maximiano, F. P. Mendonça, J. Menger, J. W. Morais, A. H. N. de Souza, J. L. P. Souza, V. da C. Tavares, J. D. do Vale, E. M. Venticinque, J. Zuanon, W. E. Magnusson, The role of environmental filtering, geographic distance and dispersal barriers in shaping the turnover of plant and animal species in Amazonia. Biodivers. Conserv. 29, 3609–3634 (2020).

104. T. R. Feldpausch, L. Banin, O. L. Phillips, T. R. Baker, S. L. Lewis, C. A. Quesada, Lloyd, Height-diameter allometry of tropical forest trees. Biogeosciences 8, 1081–1106 (2011).

105. UNEP-WCMC, IUCN, Protected Planet: The World Database on Protected Areas (WDPA) and World Database on Other Effective Area-based Conservation Measures (WD-OECM). [Preprint] (2025). http://protectedplanet.net/.

106. R Core Team, R: A Language and Environment for Statistical Computing. R Foundation for Statistical Computing [Preprint] (2024). https://www.R-project.org/.

107. R. J. Hijmans, The terra package. rspatial.org [Preprint] (2023). https://rspatial.org/pkg/terraPackage.pdf.

