## Supplementary material for "Amazon biodiversity is at risk from metal contamination due to mining activity"

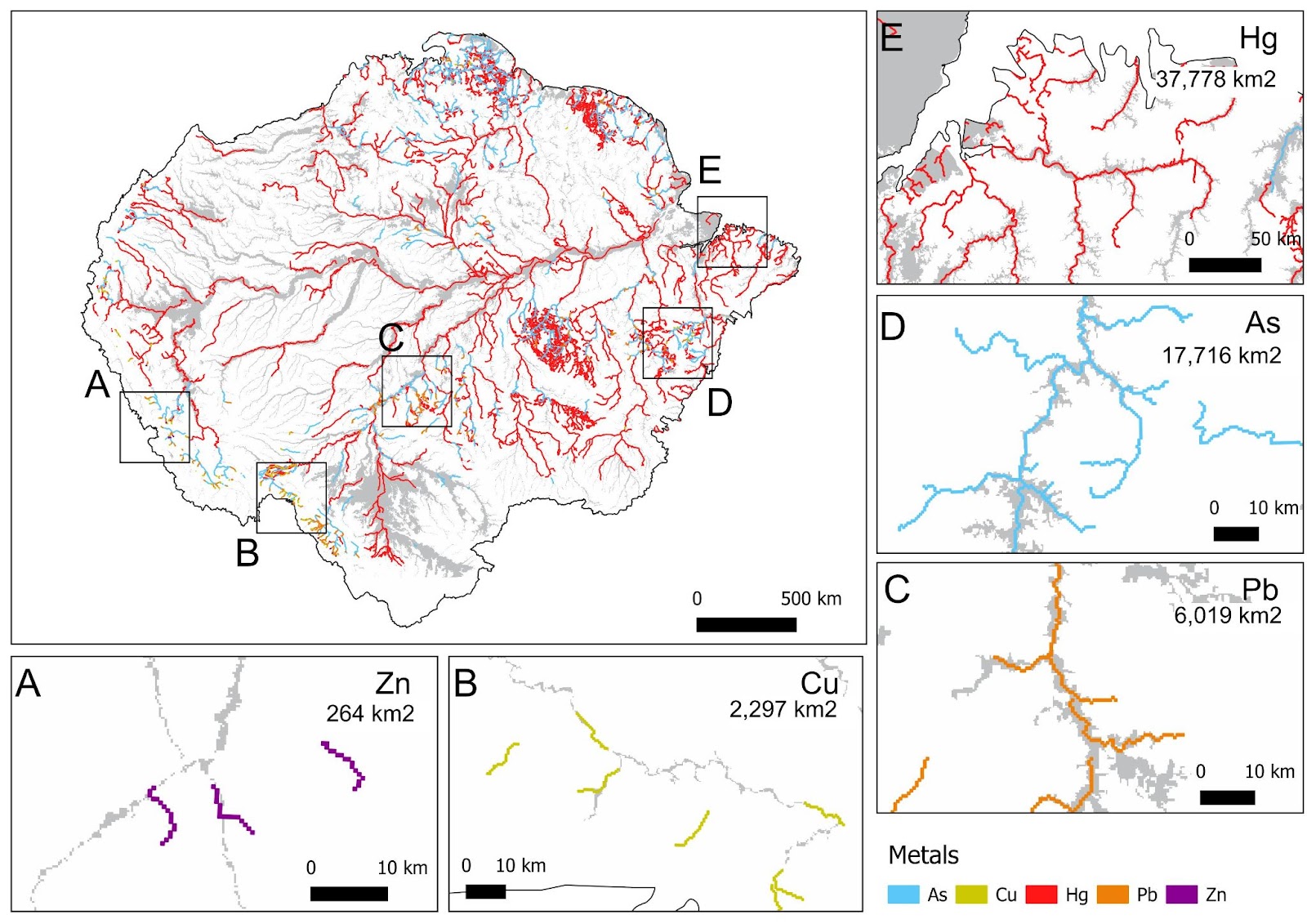

Fig. S1.

Maps of metal contamination (by mercury [Hg], arsenic [As], copper [Cu], Zinc [Zn], and lead [Pb]) along Amazonian rivers and floodplains. The extent of metal contamination is represented by the lines of different colors. Amazon main rivers as mapped by the HydroSheds project are shown in the background in grey. Contamination zones (colored lines) are those where cells (map pixels of 5 km by 5 km) downstream of mines had predicted concentrations of each metal greater than or equal to the Probable Effect Level (PEL) concentrations for freshwater sediment established by Canadian Environmental Quality Guidelines (CCME, 2024). Detailed maps A-E show examples of the distribution of the contamination along some rivers for each metal. Maps also include information on the total contamination extent of each metal in the Amazon basin.

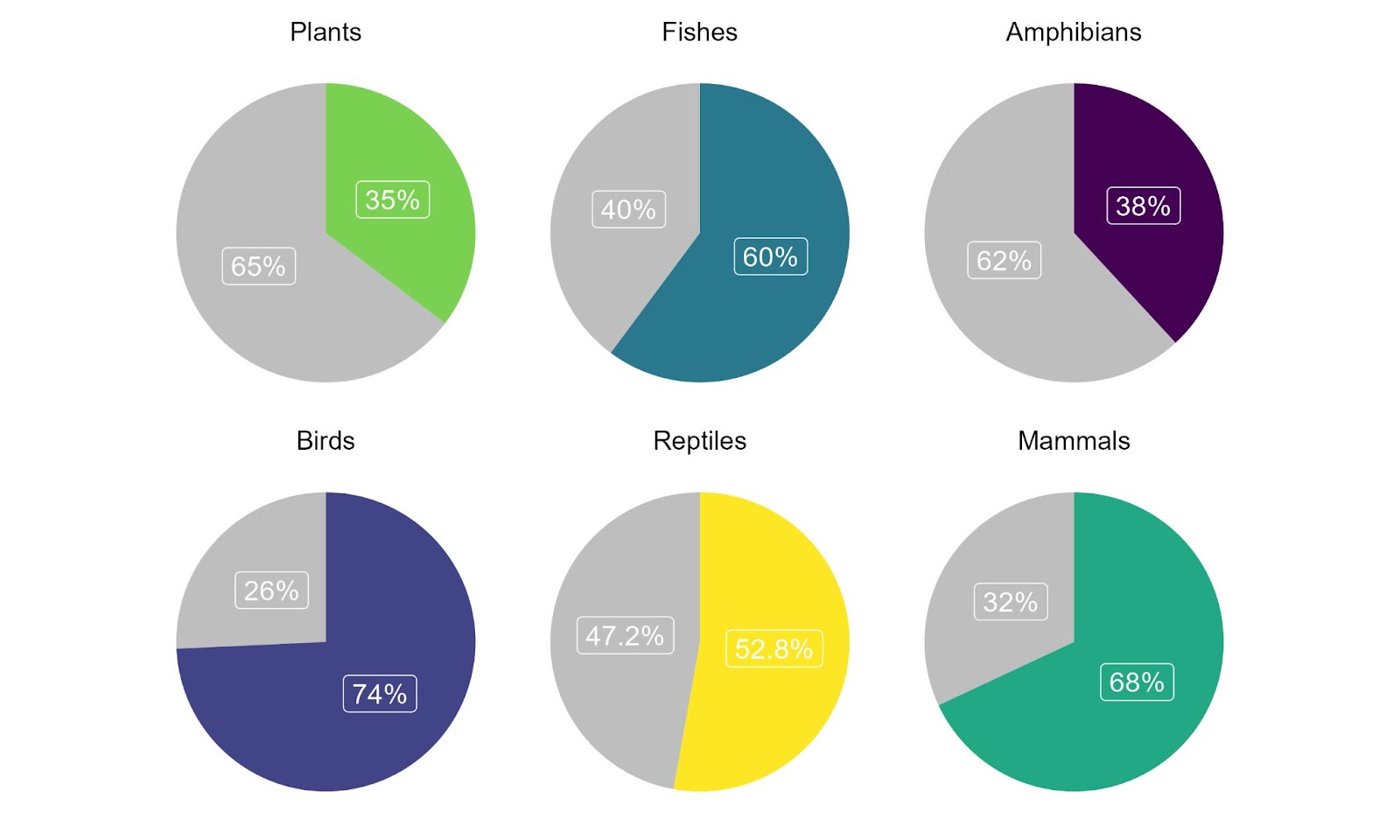
**Fig. S2.**

Pie charts show the proportion of biological species by taxa (in colours) with ranges overlapping with any of the analyzed metals (Hg, As, Pb, Zn, Cu).

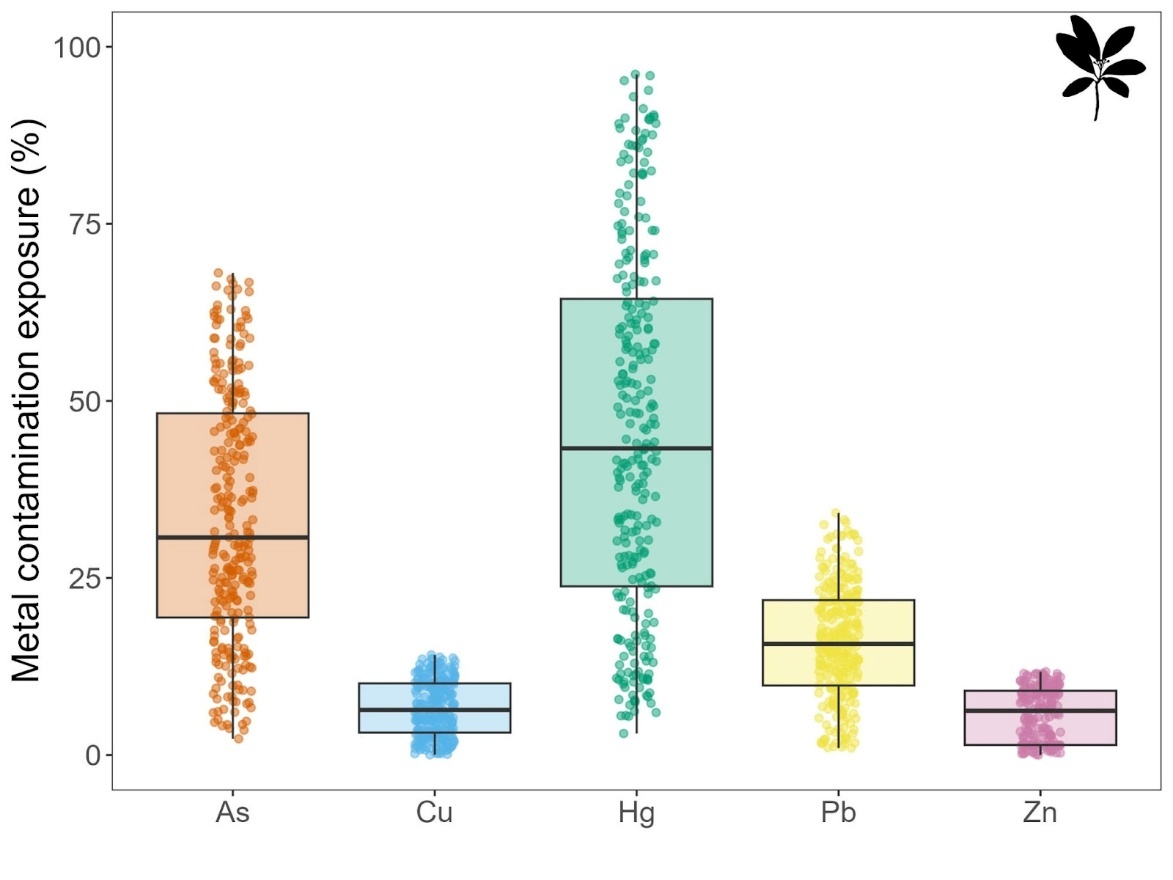

**Fig. S3.**

Ranges’ exposure of 277 riparian trees, sorted by metal contaminant. On average, 37% of riparian trees’ ranges are exposed to metal contamination (Table S1).

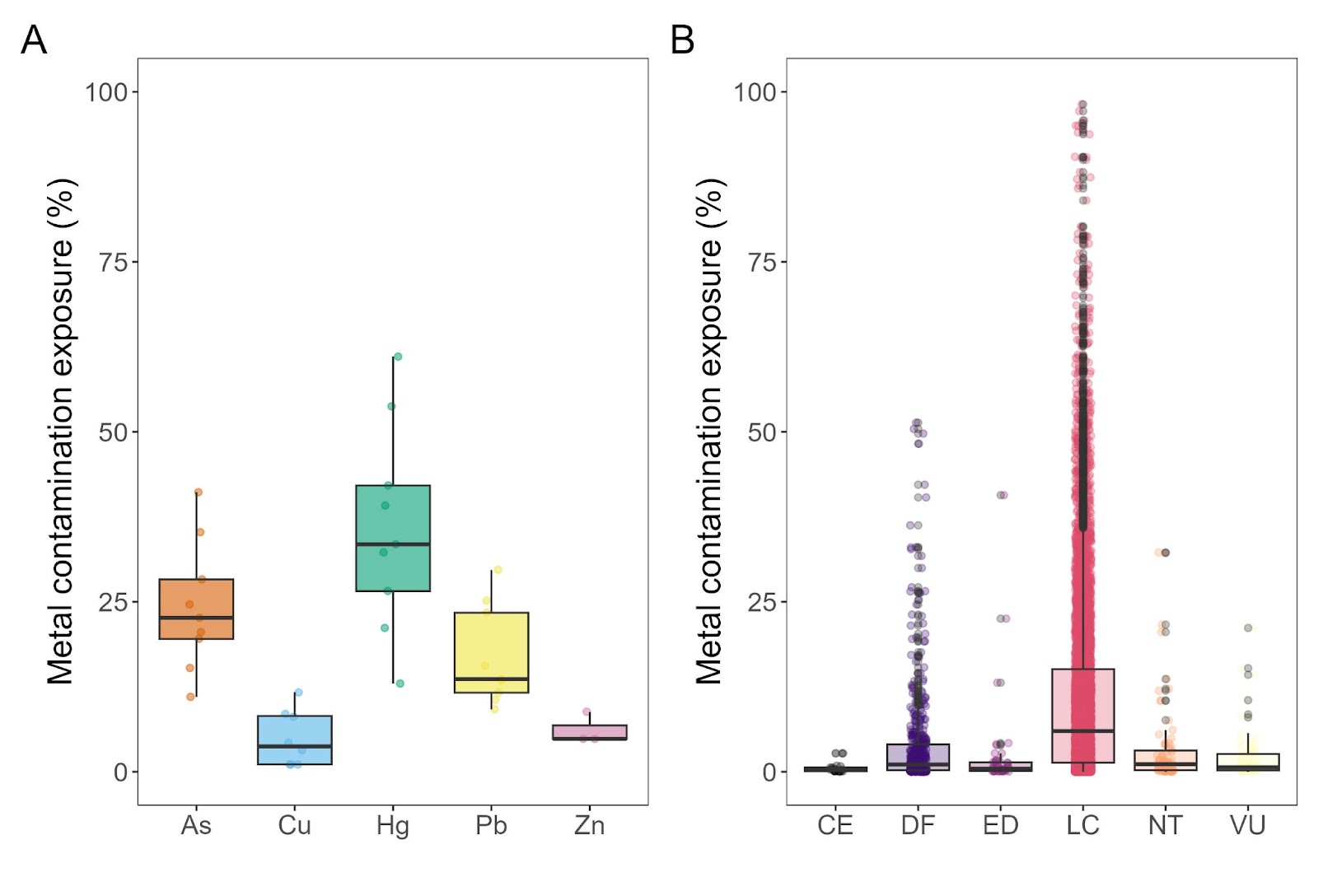
**Fig. S4.**

Ranges’ exposure of A) ten commercial fishes, sorted by metal contaminant; and B) Ranges’ exposure of all fish species divided into their current IUCN categories. The exposure is quantified as the proportion of each individual species’ ranges that overlaps with the mapped contamination extent (Figure S2). IUCN categories are: LC = Least Concern, CE = Critically Endangered, DF = Data Deficient, ED = Endangered, NT = Near Threatened, VU = Vulnerable. The ten commercial fish species are: Arapaima gigas, Brachyplatystoma rousseauxii, Colossoma macropomum, Mylossoma aureum, Mylossoma duriventre, Prochilodus nigricans, Pseudoplatystoma corruscans, Pseudoplatystoma reticulatum, Pseudoplatystoma tigrinum, and Zungaro zungaro.

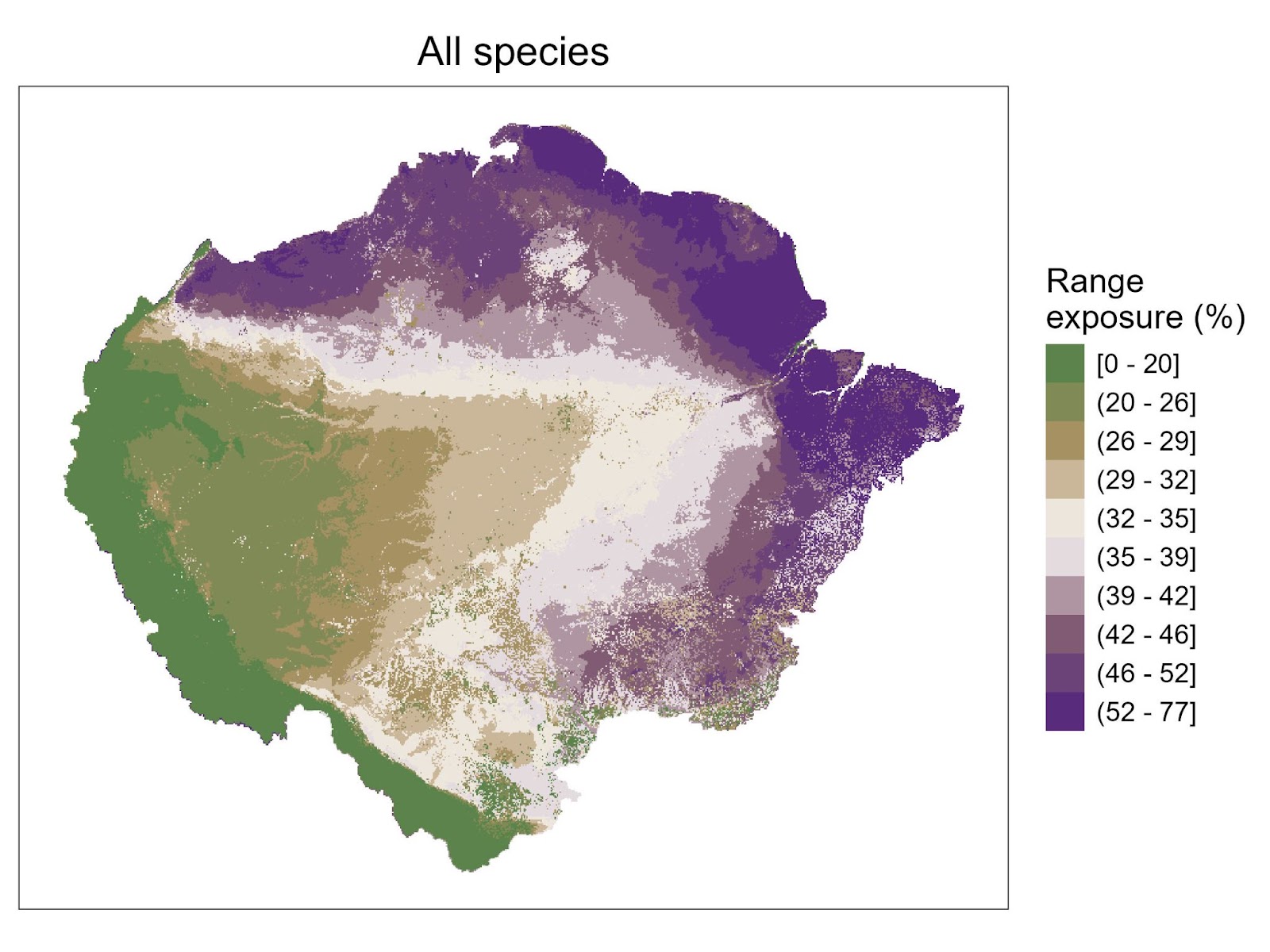

**Fig. S5**.

Map of biological species’ geographic distributions exposure for all species. geographic distributions exposure maps were made by assigning the corresponding values to their exposed areas (one value for each species geographic distributions; Figure 1), and then calculating the median values per each 5 km by 5 km grid cell.

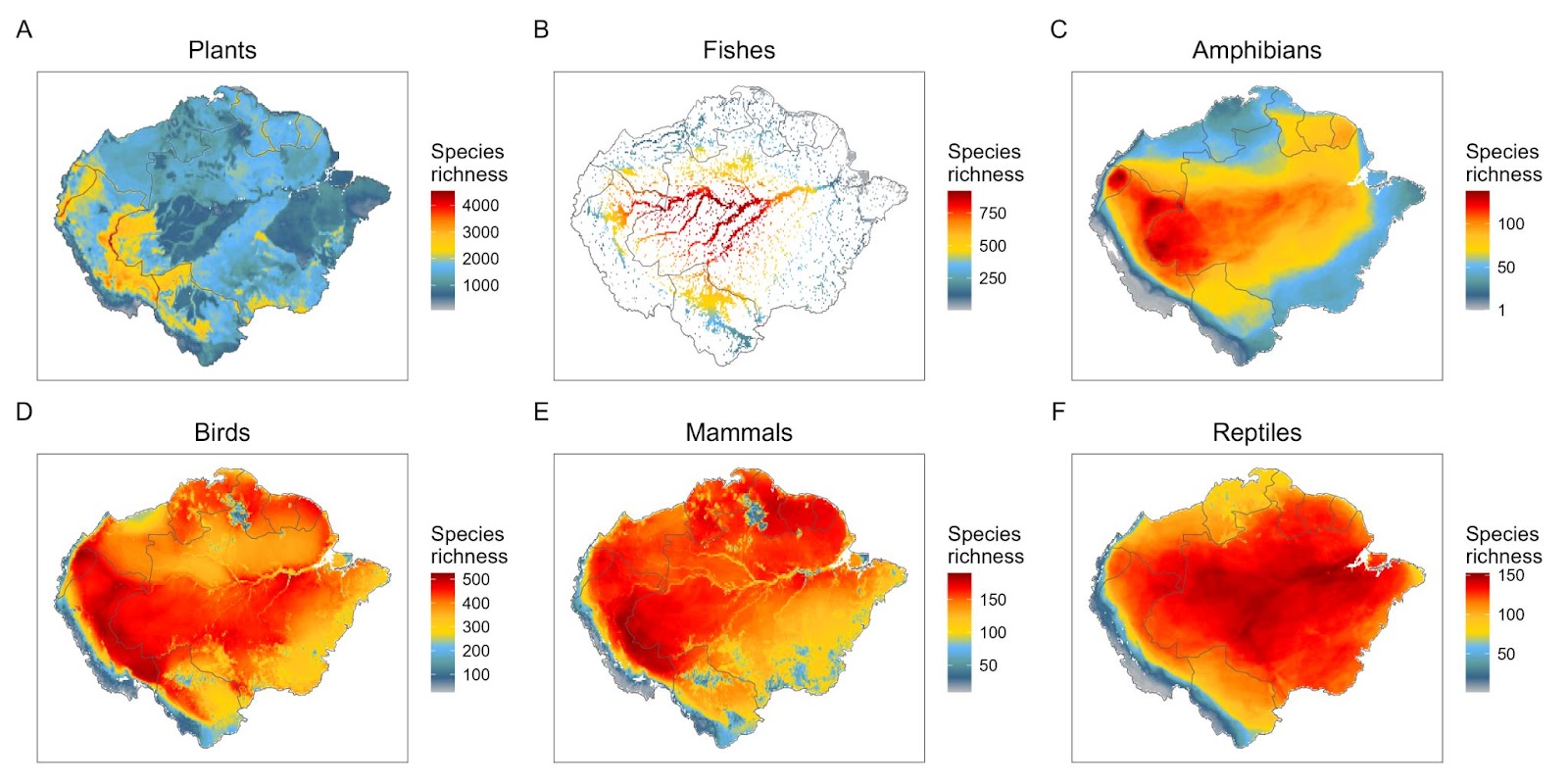

**Fig. S6.**

Maps of species richness in the Amazon basin by taxonomic groups. These maps were made by calculating the sum of overlaying species ranges per grid cell at 5 km resolution.

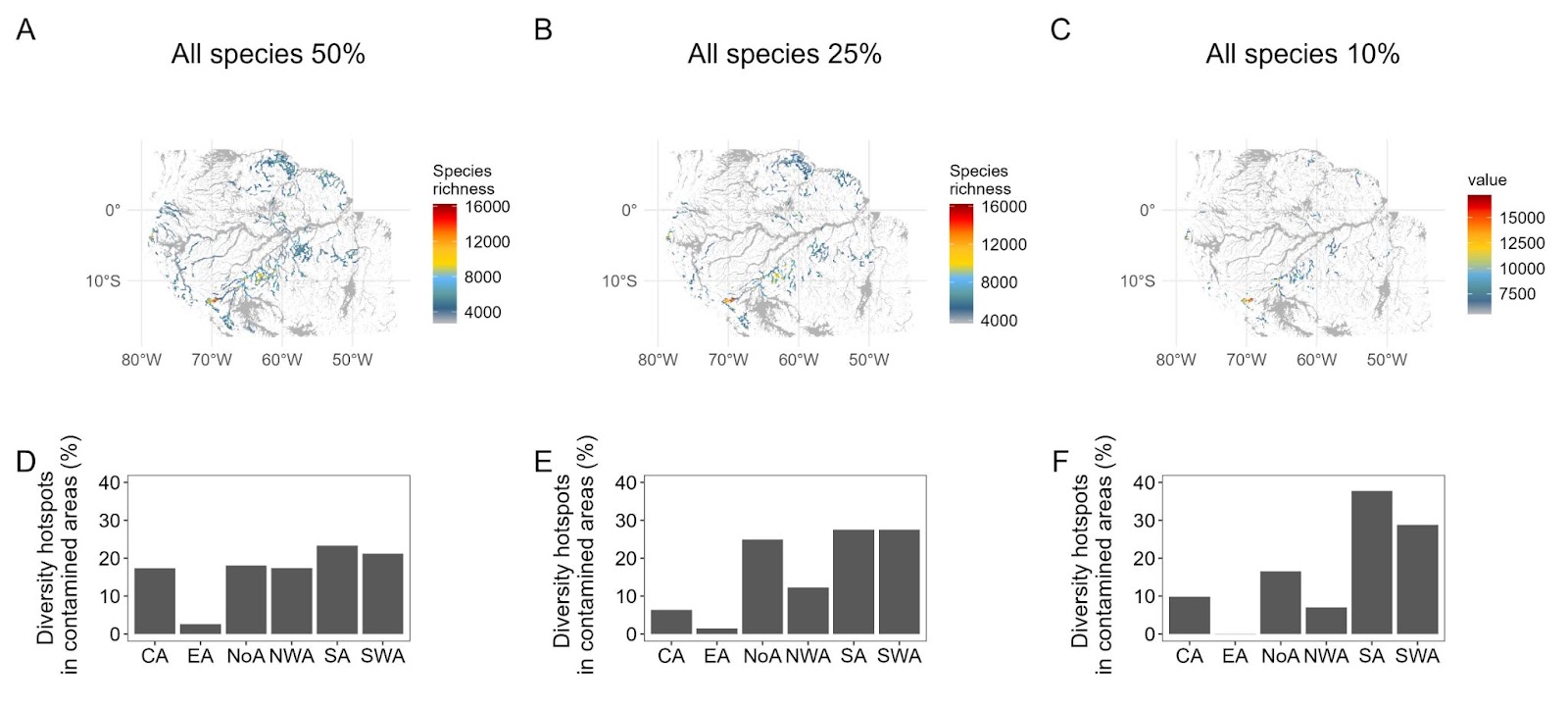

**Fig. S7.**

**(A-C)** Maps of diversity hotspots representing the overlap of species diversity with metal contamination areas resulted from the five metal contaminants (As, Cu, Hg, Pb, Zn) along the main rivers of the Amazon basin. Diversity hotspots were defined as 50, 25, and 10 % richness quantiles. **(D-F)** The proportion of species diversity overlaps with metal contamination areas in each Amazonia biogeography region. NWA = Northwestern Amazonia, SWA = Southwestern Amazonia, SA = Southern Amazonia, CA = Central Amazonia, NoA = Northern Amazonia, EA = and Eastern Amazonia. Values for the bar charts **D-F** can be found in Table S3.

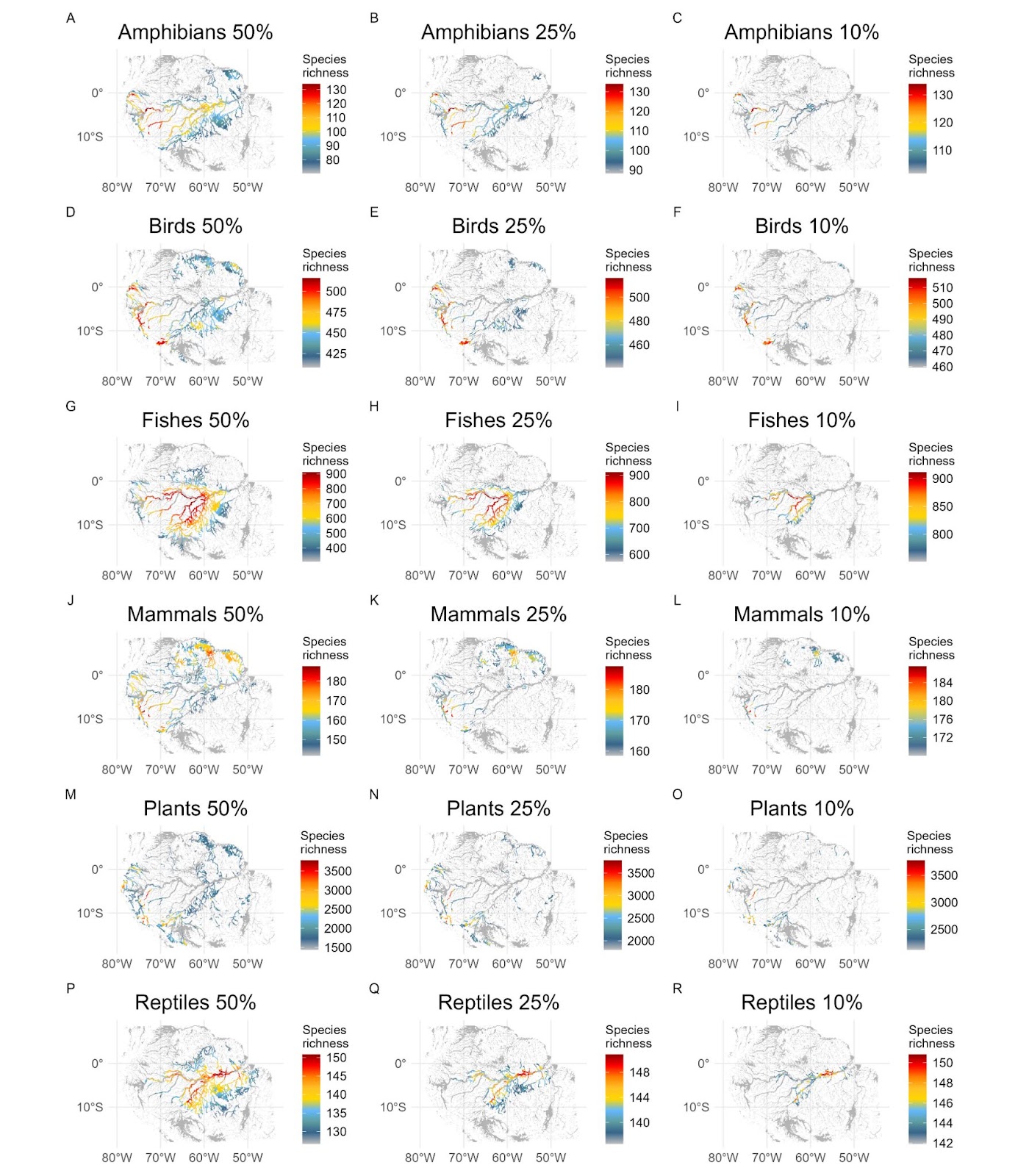

**Fig. S8.**

Hotspots of diversity of each species group that overlap with metal contamination, considering all metals. Amazon main rivers are shown in the background, in grey.

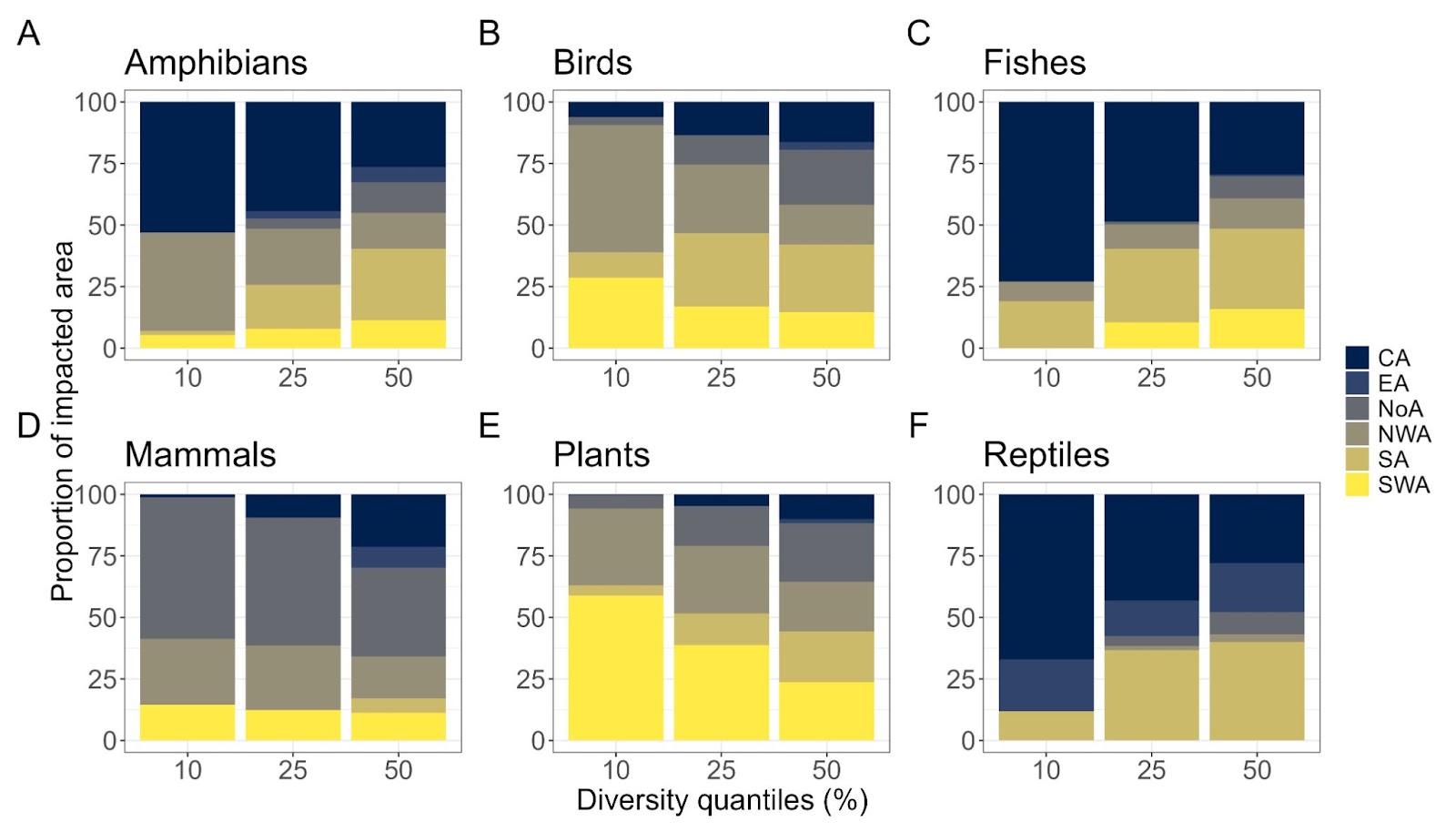

**Fig. S9.**

Proportion of exposure areas by metal contamination in each biogeochemical provinces as depicted by richness quantiles (50th , 25th and 10th percentile of species richness) and taxonomic group. NWA = Northwestern Amazonia, SWA = Southwestern Amazonia, SA = Southern Amazonia, CA = Central Amazonia, NoA = Northern Amazonia, EA = and Eastern Amazonia. Raw values can be found in Appendix 3.

Table S1. Summary of total exposure areas of each taxonomic group.

| Taxonomic group | Average exposure areas (%) | Minimum exposure areas (%) | Maximum exposure areas (%) |
| --- | --- | --- | --- |
| Amphibians | 6.80 | 0.0000 | 76.97 |
| Birds | 14.66 | 0.0000 | 80.74 |
| Fishes | 12.47 | 0.0000 | 80.22 |
| Mammals | 15.96 | 0.0000 | 76.62 |
| Plants | 2.27 | 0.0000 | 77.18 |
| Reptiles | 17.85 | 0.0000 | 79.67 |
| Riparian trees | 37.73 | 2.5732 | 78.54 |

**Table S2.** Summary of species exposition to each metal contaminant (As, Cu, Hg, Pb and Zn) across taxonomic groups. The average exposition areas indicate the overlap of individual species of each taxonomic group to the contamination reach of each metal. The number of exposed species indicates how many species of each taxonomic group has exposition above 1% (see methods). The column proportion of total taxa species indicates what is the proportion of exposed species in relation to the total number of species of each taxa.

| Metal | Taxonomic groups | Average exposition areas (%) | Exposed species | Proportion of total taxa species |
| --- | --- | --- | --- | --- |
| As | Amphibians | 1.82 | 651 | 53.71 |
| As | Birds | 3.60 | 2,059 | 87.21 |
| As | Fishes | 3.18 | 1,762 | 81.95 |
| As | Mammals | 4.15 | 603 | 81.05 |
| As | Plants | 1.91 | 15,368 | 48.39 |
| As | Reptiles | 4.47 | 446 | 67.47 |
| As | Riparian Trees | 9.24 | 276 | 99.64 |
| Cu | Amphibians | 0.05 | 405 | 33.42 |
| Cu | Birds | 0.11 | 1,745 | 73.91 |
| Cu | Fishes | 0.11 | 913 | 42.47 |
| Cu | Mammals | 0.13 | 481 | 64.65 |
| Cu | Plants | 0.10 | 10,609 | 33.40 |
| Cu | Reptiles | 0.10 | 339 | 51.29 |
| Cu | Riparian Trees | 0.24 | 270 | 97.47 |
| Hg | Amphibians | 5.76 | 686 | 56.60 |
| Hg | Birds | 11.95 | 1,964 | 83.19 |
| Hg | Fishes | 9.21 | 1,944 | 90.42 |
| Hg | Mammals | 12.87 | 606 | 81.45 |
| Hg | Plants | 0.66 | 13,370 | 42.10 |
| Hg | Reptiles | 14.52 | 475 | 71.86 |
| Hg | Riparian Trees | 26.74 | 276 | 99.64 |
| Pb | Amphibians | 0.38 | 470 | 38.78 |
| Pb | Birds | 0.61 | 1,867 | 79.08 |
| Pb | Fishes | 0.78 | 1,494 | 69.49 |
| Pb | Mammals | 0.71 | 524 | 70.43 |
| Pb | Plants | 0.48 | 11,355 | 35.75 |
| Pb | Reptiles | 0.79 | 360 | 54.46 |
| Pb | Riparian Trees | 1.51 | 276 | 99.64 |
| Zn | Amphibians | 0.01 | 99 | 8.17 |
| Zn | Birds | 0.01 | 1,133 | 47.99 |
| Zn | Fishes | 0.02 | 356 | 16.56 |
| Zn | Mammals | 0.01 | 317 | 42.61 |
| Zn | Plants | 0.02 | 5,386 | 16.96 |
| Zn | Reptiles | 0.01 | 126 | 19.06 |
| Zn | Riparian Trees | 0.02 | 201 | 72.56 |

Table S3. Percentage of overlap of the biological species richness in the 10th, 25th and 50th percentile (i.e. hotspots classes) with metal contaminated areas in each biogeographic region. NWA = Northwestern Amazonia, SWA = Southwestern Amazonia, SA = Southern Amazonia, CA = Central Amazonia, NoA = Northern Amazonia, EA = and Eastern Amazonia.

| Bio region | Hotspot class (%) | Overlap (%) |
| --- | --- | --- |
| NoA | 10 | 2.00 |
| SA | 10 | 4.58 |
| CA | 10 | 1.19 |
| EA | 10 | 0.01 |
| NWA | 10 | 0.84 |
| SWA | 10 | 3.50 |
| NoA | 25 | 7.38 |
| SA | 25 | 8.15 |
| CA | 25 | 1.89 |
| EA | 25 | 0.42 |
| NWA | 25 | 3.61 |
| SWA | 25 | 8.13 |
| NoA | 50 | 11.03 |
| SA | 50 | 14.27 |
| CA | 50 | 10.64 |
| EA | 50 | 1.57 |
| NWA | 50 | 10.70 |
| SWA | 50 | 13.00 |

**Appendix 1:** List of the biological species considered, with the associated proportion of their geographic distribution areas exposed to metal contamination.

**Appendix 2:** List of the biological species affected both by metals (this study) and fire (Feng et al. 2021).

**Appendix 3:** Percentage of overlap of the species richness in 10th, 25th and 50th percentile (i.e. hotspot classes) with mining contaminated areas in each biogeographic region.

**Appendix 4:** Maximum range exposure (%) within Indigenous territories.
